# Investigating GERMs: How Genotype, Environment, and Rhizosphere Microbiome interactions underlie heat response in maize and sorghum

**DOI:** 10.64898/2025.12.10.693489

**Authors:** Nate Korth, Isabella Borrero, Katelyn Rumley, Alex L. Woodley, Mallory J. Choudoir, Joseph L. Gage

**Affiliations:** Department of Crop and Soil Sciences, North Carolina State University, Raleigh, NC 27695; NC Plant Sciences Initiative, North Carolina State University, Raleigh, NC, 27606; Department of Plant and Microbial Biology, North Carolina State University, Raleigh, NC 27695

## Abstract

Plant resistance to heat stress can be modelled by variation attributable to the genotype, environment, the rhizosphere microbiome, and their interactions. Using this Genotype × Environment × Rhizosphere Microbiome (GERMs) model, we studied three cereal genotypes: two inbred maize lines with contrasting heat sensitivity, and a sorghum inbred that displayed moderate heat tolerance. Plants were grown under optimal and heat stressed conditions across two soil treatments. We developed a systems-level metatranscriptomics approach to examine both plant and microbial transcriptomic profiles and integrated them with microbiome compositional data and plant phenotypes. We compared our strategy to amplicon profiling and found that our metatranscriptomic strategy offers greater functional and taxonomic resolution, allowing us to characterize active microbial pathways and analyze them jointly with plant gene expression profiles within a single system. We show that the microbiome functional profile is driven by host genotype and environmental factors and can enhance plant resilience. Our analyses identified plant genes and microbial pathways consistently associated with heat tolerance and key host–microbe interactions. Specifically, we identified D-amino acid metabolism as a plausible mechanism underlying a synergistic response to heat stress. These results demonstrate that the rhizosphere microbiome is not a passive component but an active participant in plant responses to abiotic stress. This work offers a new perspective on cereal adaptation to high temperatures and underscores the utility of the GERMs framework for dissecting functional relationships among plant genotype, environment, and the rhizosphere microbiome.

## Introduction

Genotype-by-environment interactions (GxE) are a central area of research for the future of sustainable agriculture^1,2^. While it is well established that an organism’s phenotype is shaped by its genotype, the environment, and their interaction, most GxE models do not account for the host-associated microbiome. The recognition of the host and its associated microbes as a single functional unit—a holobiont— has fundamentally reshaped our understanding of biology^3,4^. Recent studies have demonstrated that GxE can significantly impact mutualism-related traits in legumes, among many other phenotypes^5,6^. Given the crucial role of the host microbiome in fitness, we echo Oyserman et al. in proposing that microbiome information be incorporated into GxE models as a distinct term^7^. The Genotype, Environment, Rhizosphere Microbiome (GERMs) interactions framework allows us to dissect these complex relationships, providing a deeper understanding of the effects of each component and their interactions on host phenotypes. In this paper, we apply this framework to identify how the rhizosphere microbiome contributes to plant adaptation to heat stress and underlies GxE mechanisms.

The GERMs framework is particularly useful for addressing the challenges posed by climate change, as rising global temperatures present an immense threat to agriculture^8^. 2024 was the warmest year on record, with global average temperatures 1.55 °C higher than the pre-industrial average^9^. High temperatures early in the growing season can increase crop growth rates, but also have damaging effects on plant maturation and grain fill^10^. Bacterial and fungal members of the rhizosphere microbiome are known to increase thermotolerance in multiple plant species, likely through the preactivation of heat-shock transcription factors and upregulation of plant secondary metabolites^11,12^. Many plants, including cereal crops such as *Zea mays L.* (maize) and *Sorghum bicolor* (sorghum), recruit specific rhizosphere microbes under heat and other abiotic stress through the production of specific root exudates^13,14^.

Cereal crops, which account for more than half of the calories consumed by humans worldwide, are threatened by climate change^15^. More than 1 billion tons of maize are produced worldwide each year, making it the world’s most-produced grain and an extremely important crop for global food and fuel production^16^. Thus, minor improvements to the sustainability of modern maize can have a massive impact. While sorghum is globally important, ranking as the fifth most widely grown cereal, it is also a close relative of maize and shares many orthologous genes^17^. Sorghum, through its human-aided adaptation across nearly all of continental Africa, also exhibits tolerance to many abiotic stresses, including heat and drought^18,19^. While maize and sorghum may share some mechanisms for resisting abiotic stress conserved across evolutionary time, mechanisms of resilience unique to sorghum could be incorporated into maize through gene editing, and insights into possible differences in sorghum-microbe interactions vs maize-microbe interactions could also prove useful. While genetic mechanisms are often conserved, mechanisms of microbiome recruitment and signaling based on microbial function remain obscured by methods that focus solely on microbial community composition.

While the field has made strides in understanding plant-microbe interactions under climate stress, much of the existing literature focuses on a single crop species and a single plant genotype or variety within that species. The few studies examining the microbiomes of multiple plant species or genotypes under abiotic stress conditions rely on microbiome amplicon sequencing^7,20,21^. Research on the human microbiome suggests that while the microbial composition of a given body site varies significantly between individuals, the microbial functional profile, as quantified by transcriptomics or proteomics, remains remarkably conserved^22^. While there is a lack of overlapping compositional and functional datasets from the plant rhizosphere microbiome, compositional data from multiple environments show high levels of variation between field sites (environments)^23^. Despite strong environmental drivers of rhizosphere composition, inferred functional enrichment showed far less variation in a phenomenon similar to that observed in humans^24^. To better understand the functional relationships among maize, the environment, and the plant-associated microbiome, we employed a systems-level metatranscriptomic approach. This method derives functional information from multiple kingdoms of life from a single preparation of RNA extraction and metatranscriptomic sequencing. We designed a bioinformatic workflow to separate mRNA from plants, bacteria, and fungi, and to explore mechanisms underlying genetically controlled maize microbiome-associated phenotypes.

In this study, we investigate how plant and rhizosphere microbial gene expression jointly respond to temperature stress and influence plant performance. By integrating transcriptomic data from both host and microbes, we identify plant genes and microbial pathways associated with heat stress resilience. Using co-expression networks, multivariate ordination, and machine learning, we pinpoint plant-microbe interactions consistently linked to adaptive traits. This integrative approach provides a systems-level view of the molecular mechanisms underlying plant resilience to abiotic stress.

## Materials and Methods

### Soil preparation

Field soil was collected from North Carolina State University’s Lower Coastal Plain Research Station in Kingston, North Carolina (35.378305, -77.559656) from fields with maize and wheat rotations, with the most recent rotation being maize, and kept in isolation for 60 days. The field soil is classified as Goldsboro loamy sand with a mean particle diameter (D_50_) of 0.16 mm and a plasticity index of 8. Complete soil chemistry is available in Supplemental Table 1. A portion of the soil was then subjected to two rounds of autoclaving at 212°C for 60 minutes. To alleviate compaction and ease root extraction, the potting media was formulated with 60% field soil (autoclaved or unautoclaved), 30% sterilized calcined clay (Turface™), and 10% sterile perlite (Miracle-Gro ®). The soil was mixed in 6 L batches, including 30 mL of multicote (14-14-16) nutrient solution, and added to 4-inch pots. Each pot was topped with a 1 cm layer of the autoclaved soil to reduce weed growth. Pots were then watered with RO water for one week prior to the start of the experiment to remove any weed seedlings that persisted.

### Genotype Selection

Two tropical, inbred maize genotypes, a heat-tolerant genotype, CML52 (PI 595561), and a heat-susceptible genotype, CML103 (PI 690319), were obtained from a collection of the maize 282 panel grown to increase seed in Puerto Vallarta, Mexico, in 2022-2023^25,26^. A moderately tolerant genotype of sorghum, SAP-166 (PI 597964), was obtained from the North Central Regional Plant Introduction Station (NCRPIS) in Ames, Iowa, in 2022.

### Experimental Design and Growth Chamber Conditions

Each genotype was replicated four times in field and autoclaved soil at two temperature conditions. Autoclaved soils served as a microbiome-depleted condition. Bulk field and autoclaved soil in pots without plants at both temperatures were included in triplicate as controls. Pots were randomized in a complete block design across two growth chambers: one at normal growth conditions (28°C -day and 20°C -night) and the other under heat stress conditions (38°C -day and 28°C -night). The light intensity of both growth chambers was set to 320 μmol/(s · m²)^27^.

### Seedling Germination

Seeds were sterilized by soaking in a 10% sodium hypochlorite solution with one drop of Tween® 20 on a rotating mixer for 20 minutes. Seeds were then triple-rinsed with sterile H_2_O before soaking in 70% EtOH for 2 minutes. To finish, seeds were triple-rinsed with sterile H_2_O. Filter paper was wet and layered on top of plastic wrap and wax paper. Seeds were then all placed one inch apart, two inches below the top of the filter paper with the germ pointed downward. A second piece of damp filter paper and wax paper were layered on top, and all five layers were then loosely rolled and placed into a large beaker with water, ensuring that seeds were not submerged. The beaker was then placed in a growth chamber set to 28_°_C/20_°_C. After two days, the rolls were checked for germination and transplanted if germination had occurred. Pots were watered to saturation at the time of transplanting and then as needed until sampling.

### Plant Sampling for RNA and DNA Extractions

The day of sampling was decided based on when the majority of the plants reached V3-V4 on the plant growth stage scale^28^. The morning of sampling, plant growth stage was noted, a photo of each pot was quickly taken, and plants were rated on a 0-4 categorical scale based on plant size and leaf senescence, where zero means the plant showed no sign of heat stress and four means that the plant showed extreme stress and was withered (Figure 1).

**Figure 1.**
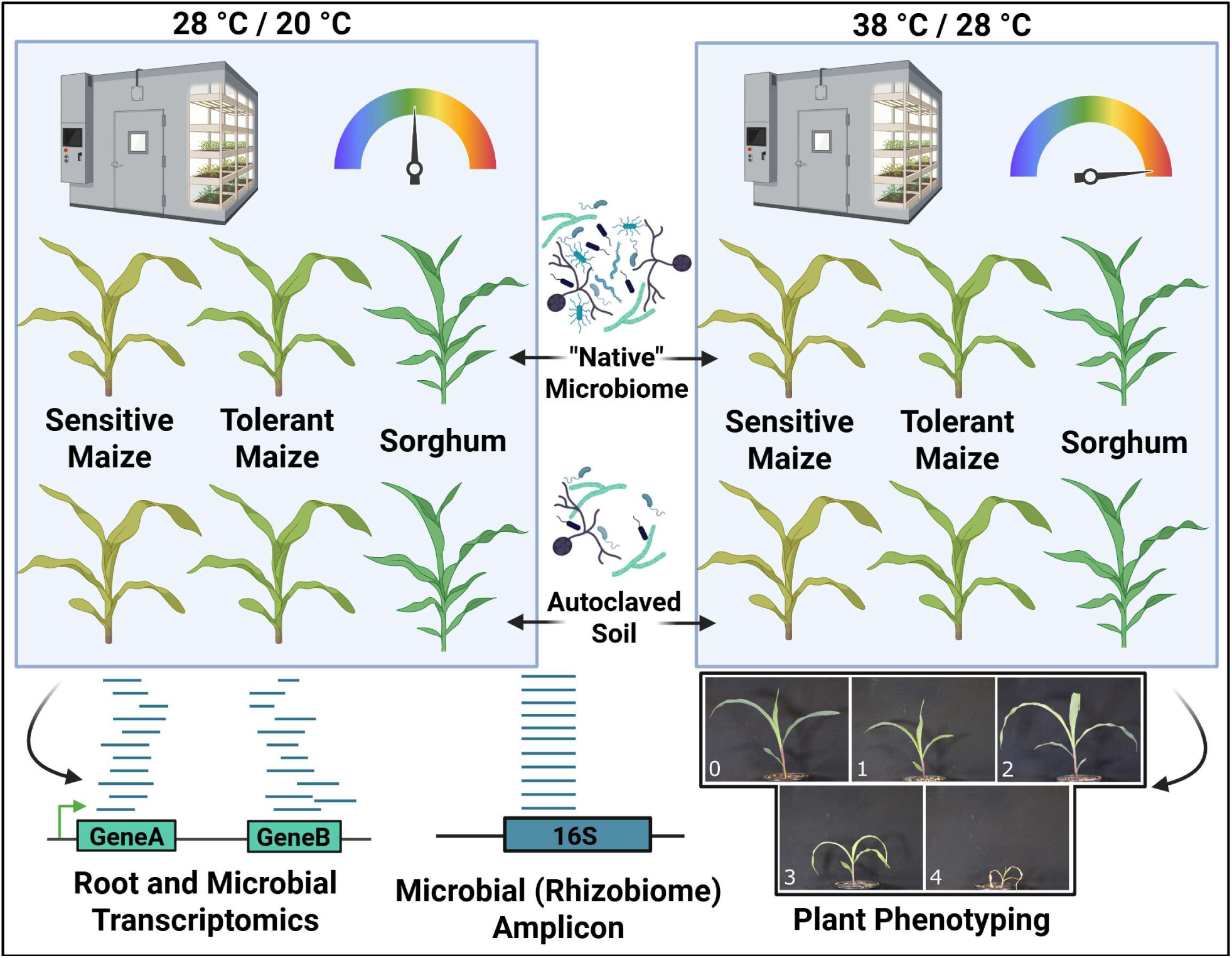
Graphical representation of methods. Three genotypes — a heat-resistant maize, a heat-susceptible maize, and a sorghum variety — were grown to the V4 stage in growth chambers under optimal conditions or subjected to heat stress. Plants were grown in soil containing a complex community of microbes, or in the same soil that had been autoclaved. Total RNA from roots and root-associated microbes was sequenced, along with the 16S rRNA amplicon from both DNA and RNA. Plants were assigned a qualitative heat stress score based on size and leaf senescence (Bottom Right), as well as quantitative metrics, including biomass and root architecture.

Roots were sampled in sets of four plants, with two people working simultaneously. To remove plants, pots were gently squeezed to loosen soil and inverted onto trays.

Excess soil was manually massaged from the roots, which were then gently shaken to remove loose particles. Large particulates were removed, and shoots were clipped at the base and set aside for biomass measurements.

Cleaned roots were sonicated to loosen adhered soil, microbes, and the outer layer of plant cells, yielding a mixture of microbial and plant nucleic acids in each sample. Roots were immediately fully submerged into pre-chilled 50 mL tubes containing 25 mL of 1× phosphate-buffered saline (PBS; pH 7.4). Tubes were inverted five times, sonicated for 20 s, inverted, and sonicated for an additional 20 s, followed by five more inversions. Roots were then removed, leaving soil in suspension. Samples were centrifuged at 2,500 × g for 1 min at 4 °C, and the supernatant was discarded. Pellets were flash-frozen in liquid nitrogen and stored at −80 °C until nucleic acid extraction.

### Plant phenotyping

In addition to the categorical heat stress score, biomass from cleaned roots and shoots was collected. Roots were then given an additional wash and scanned using a WinRHIZO root scanner (Regent Instruments Inc., Québec, Canada) equipped with the WinRHIZO Pro software (version 2022b). Roots were carefully spread in a water-filled, transparent tray to minimize overlap, then scanned at high resolution. Images were analyzed in WinRHIZO/XLRhizo using the manufacturer’s default calibration and classification settings to quantify root system architecture traits. A range of root phenotypes were extracted, including total root length, average diameter, surface area, volume, and topological parameters (e.g., tips, forks, crossings). The complete list of phenotypes measured, along with their corresponding values, is included in Supplemental Table 2 (Metadata).

### DNA and RNA Extractions

RNA and DNA were extracted from the frozen soil samples using the Qiagen PowerSoil DNA Kit (12866-25) and the RNeasy PowerSoil Total RNA Kit (12867-25), respectively.

### Amplicon Sequencing (DNA)

We surveyed 16S rRNA genes from DNA and RNA to profile whole communities (active and dormant taxa) and active fraction, respectively. Extracted DNA was sent to the North Carolina State University Genomic Science Laboratory for Illumina 16S/ITS Amplicon library preparation and sequencing with 16S primers 322F-A (ACGGHCCARACTCCTACGGAA) and 796R (CTACCMGGGTATCTAATCCKG) designed to amplify microbial 16S from plant tissue selectively^29^.

### Amplicon Sequencing (RNA)

To create a cDNA library, remaining DNA was removed from RNA samples using the TURBO DNA-free Kit (#AM1907; Invitrogen) DNase Kit. DNA-free RNA samples were then reverse-transcribed into cDNA using the SuperScript III Reverse Transcriptase Kit (#18080044; Invitrogen) following the standard protocol. The gene-specific primers used were the 515F (sequence: GTGCCAGCMGCCGCGGTA) and 806R primer (sequence: GGACTACHVGGGTWTCTAAT) for amplification of the V4 region of the bacterial and archaeal 16S rRNA gene, as provided by the Earth Microbiome Project^30^. The cDNA served as the template for PCR amplification: each 25 µL reaction volume contained 12.5 µL of HiFi Master Mix (M049S, New England BioLabs), 10 µL of PCR-grade H₂O, 0.5 µL of 10 µM 806R reverse primer, 1 µL of 5 µM barcoded 515F forward primer, and 1 µL of cDNA template. The following thermocycler conditions were used: an initial denaturation at 98 °C for 3 minutes; followed by 35 cycles of denaturation at 98 °C for 30 seconds, annealing at 55 °C for 30 seconds, and extension at 72 °C for 30 seconds; with a final extension at 72 °C for 7 minutes, and a final hold at 10 °C. Technical replicates were combined before amplicon concentrations were determined using the PicoGreen dsDNA Assay Kit (#254406, Invitrogen), performed on the Tecan Infinite® 200 PRO M Plex microplate reader (Männedorf, Switzerland). Samples were normalized, pooled and sequenced on a MiSeq i100 Benchtop Sequencer by Illumina (San Diego, California) using the custom primer protocol with the NextSeq 1000/2000 XLEAP-SBS Read & Index Primer Kit (#20112856; Illumina), PhiX Control v3 (FC-110-3001; Illumina) spiked in at 30%, and the MiSeq i100 Series 25M Reagent Kit (300 cycles, #20126568; Illumina). FASTQ files were generated on the Illumina secondary analysis platform BaseSpace using the DRAGEN BCL Convert 4.3.13 application.

### RNA library prep/sequencing

Extracted RNA was sent to Novogene (Sacramento, CA, USA) for library preparation and sequencing. Ribosomal RNA (rRNA) was removed using a proprietary rRNA depletion kit (Novogene). Libraries were constructed with the ABclonal® Fast RNA-seq Library Prep Kit V2 for Illumina (ABclonal, Woburn, MA, USA) according to the manufacturer’s protocol, using both non-directional (default) and directional workflows. Libraries were sequenced on the Illumina NovaSeq X Plus platform (Illumina, San Diego, CA, USA).

### Bioinformatic Workflow -Amplicon

For both 16S amplicon sequences from DNA and RNA, demultiplexed sequence data were processed in *RStudio*, following the *DADA2* v1.34.0 pipeline to infer amplicon sequence variants (ASVs). Based on the quality score, the 16S reads from the DNA forward and reverse strands were truncated to 300 and 285 nucleotides, respectively. The 16S from RNA forward and reverse reads were both truncated at position 150. Forward and reverse reads were merged, chimeras were removed, and taxonomy levels were assigned using the SILVA NR99 16S rRNA To Species Training Database v138.2^31,32^.

Reads mapping to mitochondria or chloroplasts were filtered, and taxa present in fewer than two samples or with fewer than 10 reads in at least two samples were omitted to remove low-abundance and rare taxa. Alpha diversity metrics were computed using the phyloseq package^33^. As above, read counts were converted to relative abundance, and the center log-ratio (CLR) was computed. The CLR-transformed data were used to generate a Euclidean distance matrix for beta-diversity estimation using the ‘phyloseq’ R package. Differences between samples in the distance matrix explained by temperature, soil, and genotype were tested for statistical significance using a permutational multivariate analysis of variance (PERMANOVA) in the *Vegan* v2.6-10 R package, with 999 permutations^34^.

To compare DNA versus RNA amplicon sequencing, samples were rarified to 50K reads using *the phyloseq package*. A mean value for each taxon was calculated for each level of Genotype at a given level of Temperature and Soil Type. Spearman correlations were calculated for each comparison of taxa abundances and standard deviation. A one-sided t-test was used to determine whether the slope was significantly less than the expected value (1).

### Bioinformatic Workflow -Metatranscriptome

Adapters and low-quality reads were trimmed from the raw FASTQ files using *Trimmomatic* v0.39^35^. The *HISAT2* v2.2.1 aligner was used to filter reads that aligned with the human genome^36,37^. Any sequenced ribosomal RNA (rRNA) was identified and removed using a deep learning method implemented in *Ribodetector* v0.3.1^38^. After preprocessing, filtered sequences were aligned to the B73 v5 (maize) or BTx623 v3 (Sorghum) using *HISAT2*^39,40^. Aligned reads were assigned a feature by the featureCounts tool from *SubRead* v2.0.6^41^. Orthologous genes between the references B73 and BTx623 were identified using OrthoFinder v2.5.5, and gene counts were assigned to orthogroups using a custom R script^42^. Lowly expressed genes were removed (<75 reads in < 4 samples) prior to differential expression analysis of maize or sorghum genes conducted using *DESeq2* v1.48.1 in R v4.5.0 using the model Expression = Genotype + Temperature + Soil + Soil:Temperature^43,44^. GO enrichment analysis was performed using the Ensembl Plants database and the *biomaRt* v2.64.0 and *TopGO* v2.60.1 R packages^45–47^.

Microbiome composition was inferred from reads that did not align to the maize genome by comparing sequences to the RefSeq database using *Kraken2* v2.14^48,49^. A KEGG function was assigned to each read using the *Diamond* v2.1.8 tool and the *EggNOG* v1.12 mapper, and the *EggNOG* v5.0 database, including fungal, bacterial, and archaeal orthologs^50–52^. Microbial genes annotated via EggNOG were filtered to remove low-confidence genes (e-value < 1e-6), then collapsed into pathways by EC number and assigned functional annotations from the KEGG database using the *KEGGREST* v1.42.0 and *tidyverse* v2.0.0 packages in R^53,54^.

Low-abundance microbial genes were filtered prior to analysis, retaining only those with at least 250 reads in three or more samples. Because the number of microbial transcripts captured in each library likely represents only a subset of the total root transcriptome, expression values were normalized to relative abundance to account for differences in sequencing depth and microbial representation among samples. Read counts were subsequently transformed using the centered log-ratio (CLR) method implemented in the *Compositions* R package (v2.0-8) to address the compositional nature of the data^55^. Principal component analysis (PCA) was performed on CLR-transformed data to visualize overall transcriptional profiles across samples. Samples were grouped by genotype and soil type (field or autoclaved), and differential expression analysis was conducted using *ALDEx2* (v1.40.0) to identify microbial genes whose expression differed in response to temperature^56^. *ALDEx2* was selected because it explicitly models the compositional structure and sampling variability inherent to microbial data, making it well-suited for complex microbiome metatranscriptomes^57^. For pathway-level analyses, a more comprehensive model (*Expression ∼ Genotype + Temperature + Soil + Soil:Temperature*) was employed in *ALDEx2* to evaluate interactive effects among experimental factors.

### Bioinformatic Workflow -Coexpression

To assess coexpression, we analyzed various features of the plant and microbiome transcriptome. The following analyses were conducted in parallel using microbiome transcription data at the gene level and collapsed KEGG pathways. A Mantel test was used to examine the relatedness of plant and microbiome expression profiles by computing distance matrices of plant (Median-to-ratio transformed) and microbial (CLR transformed) transcript abundance datasets with the Vegan v2.6-10 package’s mantel command, using a Spearman correlation and 5000 permutations. PERMANOVA tests were conducted for both the plant and microbial distance matrices to evaluate the contribution of genotype, soil type, and temperature to variation in gene expression. We conducted a differential correlation analysis to compare pairwise co-expression patterns at 28°C and 38°C. Pairwise Spearman correlations were then computed between all plant genes and microbial features within each temperature condition, excluding intra-domain correlations (i.e., only plant–microbe pairs were tested). For each pair of plant and microbial genes, Fisher’s *z*-transformation was applied to correlation coefficients at both temperatures, and the difference between *z*-scores was used to quantify changes in association strength between 28°C and 38°C. The standard error of the difference was calculated assuming independent sampling, and a two-tailed *z*-test was used to determine statistical significance. False discovery rate (FDR) correction was applied using the Benjamini–Hochberg method, and significant differential correlations were defined as those with FDR < 0.05 and |Δ*z*| > 1.

To identify plant genes and microbial pathways most strongly associated with key traits, we applied two complementary predictive modeling approaches: elastic net regression and random forest. Prior to modeling, plant and microbial abundance matrices were scaled, and features with weak univariate correlations (Spearman’s Rho > 0.05) with the response variable were filtered out to reduce dimensionality. For elastic net regression, models were fit separately for biomass and heat stress using a 50:50 mixture of L1 and L2 penalties (α = 0.5) using the *glmnet* v4.1-10 R package^58^. Ten-fold cross-validation was performed to select the optimal regularization parameter (λ), and nonzero coefficients were extracted to identify features contributing most strongly to trait variation. Features were annotated by type (plant gene or microbial pathway), and separate columns were retained for biomass and heat stress-associated coefficients. Random forest models were fit to the same filtered datasets using 2000 trees and 10-fold cross-validation in the *randomForest* v4.7-1.2 R package^59^. Variable importance was calculated for each feature, and the top 25 plant genes and microbial pathways were extracted for each trait.

Results were summarized to identify features consistently represented across modeling approaches, enabling the integration of co-expression analysis methods to highlight plant-microbe interactions linked to heat response phenotypes.

All data visualization was conducted in R using the *ggplot2* v3.5.1 and *ggpubr* v0.6.0 packages^60,61^.

## Results / Discussion

### Plant Phenotype Analysis

The plant genotypes responded as expected to heat stress, with survival rates> 90% across genotypes. Plants that died before the end of the experiment were assigned a heat stress value of 4 and excluded from RNA sequencing. The heat-tolerant maize genotype, CML52, exhibited the lowest heat stress scores at 38 °C, while the heat-susceptible maize genotype, CML103, showed the highest, and the sorghum genotype displayed an intermediate phenotype (Figure 2A). To verify the heat stress score, we treated it as a numeric variable and found it to be negatively correlated with biomass and most root phenotypes (Figure 2B). We observed significant effects of both soil type and temperature on the heat stress score (Fig. 2C). While CML52’s heat response remained consistent across soil types, its shoot biomass was significantly higher in autoclaved soil than in field soil (Wilcoxon, p=0.017).

**Figure 2.**
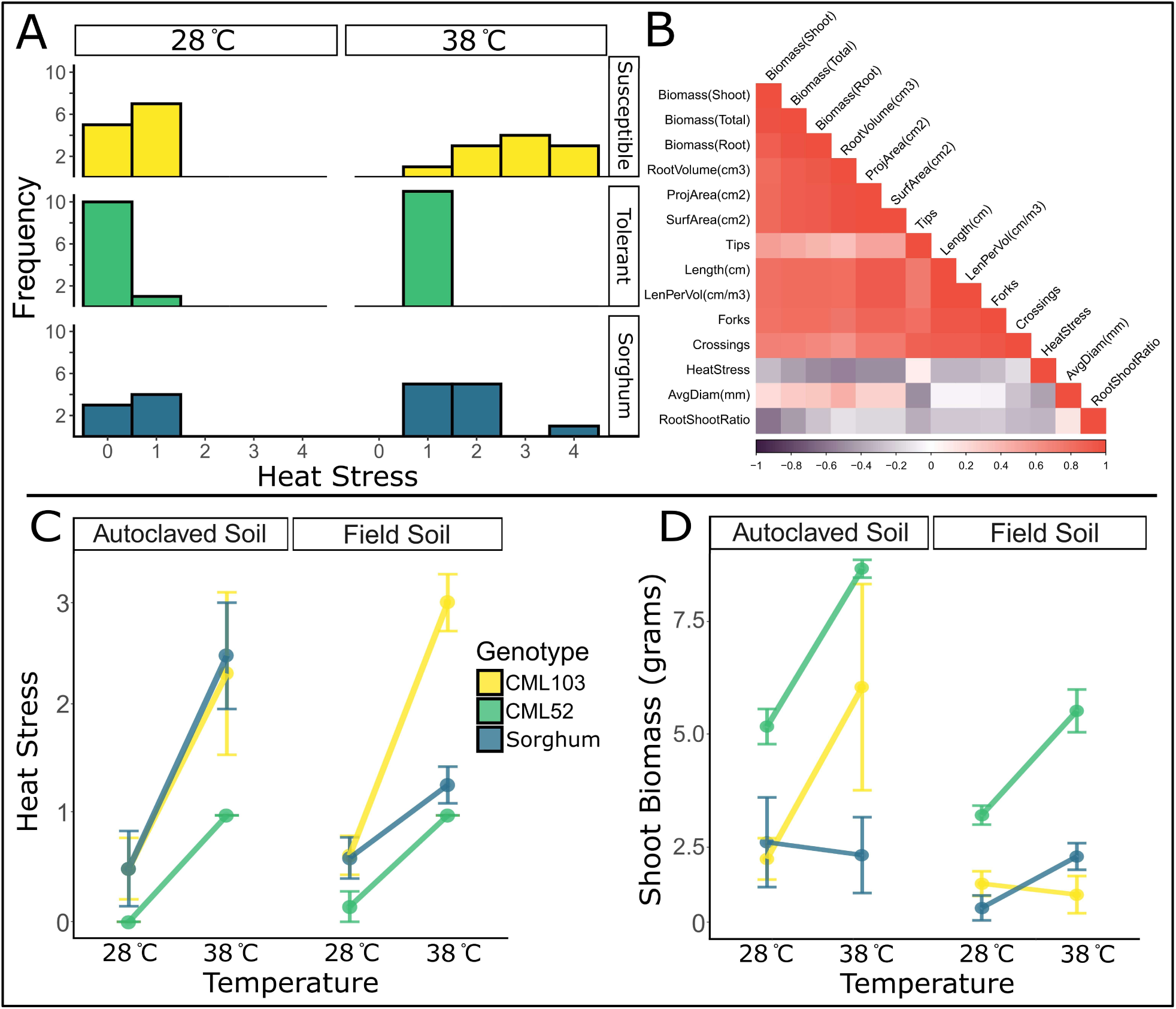
Plant phenotype analysis. The categorical heat stress score was distributed as expected across genotypes (A). The heat stress score was negatively correlated with plant biomass and root phenotypes, which were highly correlated with each other (B; Spearman’s Rho). Both soil treatment and temperature had a significant impact on heat stress (C). Raised temperatures resulted in increased biomass, provided that 38 °C wasn’t lethal to the plant (D).

In field soil, CML103’s heat stress score at 38 °C was higher than in autoclaved soil (Fig. 2C). Interestingly, the sorghum genotype exhibited the opposite trend, with a higher heat stress score in autoclaved soil at 38 °C compared to field soil. The effect of soil type on shoot biomass and root volume also showed contrasting trends between CML103 and the sorghum genotype, where sorghum appears to be more protected from heat stress in field soil, and CML103 is more protected in autoclaved soil (Fig. 2D and supplemental figure 1A). Unlike the qualitative heat stress score and above- or below-ground biomass, the root-to-shoot ratio consistently decreased in response to the high-temperature treatment, similarly across genotypes in both soil treatments, consistent with the literature (Supplemental Figure 1B)^62^. We used these four phenotypes (quantitative heat stress score, shoot biomass, root volume, and root-to-shoot ratio) as biomarkers of heat stress.

There is a high degree of variation in CML103 phenotypes in autoclaved soil and at high temperatures (Figure 2C and D). However, this observation is not unexpected; many microbiome treatments in otherwise identical samples result in varied phenotypes, referred to as the responder and non-responder phenomena in animal systems^63^. Though attention is not typically drawn to unexplained variation, its presence in plant studies underscores that the effect of microbiome perturbation is not always uniform^64,65^. To address the issue of responder/non-responder, most downstream analyses were conducted to identify plant genes and microbial composition or function that impact the plant phenotype, agnostically of the experimental design.

The results demonstrate that the magnitude and direction of a plant’s response to heat stress depend not only on the plant’s genotype, abiotic environment, and the GxE interaction, but also on the underlying context of the rhizosphere microbiome. These findings underscore the power of the GERMs framework in elucidating complex plant phenotypes. Identifying these complex interactions is the first step in uncovering the microbiome-mediated mechanisms by which plants adapt to stress.

### System-level metatranscriptomic sequence assignment

Reads from each sample were parsed into those belonging to maize/sorghum, bacteria, or fungi after removal of human reads and ribosomal RNA (rRNA). Of an average read count of 57.7M (range: 41.7M -81.1M), ∼3.4M (range: 0.62M -9.5M) reads were aligned to the maize B73 genome in maize samples, while ∼3.2M (range: 1.4M -7.1M) reads were aligned to the sorghum Tx623 genome in sorghum samples. The remaining reads were assumed to be microbial and were assigned a function and binned into those belonging to bacteria (average: 22.7M, range: 9.4M-53.14M) or fungi (average: 2.1M, range: 0.31M-5.7M) using the EggNOG database (e-value < 1e-6). Reads were given a lower taxonomic assignment based on sequence similarity to genomes in the RefSeq database (Supplemental Figure 2). The complete list of per-sample read counts is available in Supplemental Table 3.

Mapped read counts to maize and sorghum features varied widely and were below optimal coverage. Improved root cleaning prior to sonication could increase the microbial-to-plant read ratio, and higher sequencing depth per sample would further mitigate this limitation. Less stringent alignment might recover more plant features, but risks misassigning fungal reads to plants. Low plant read counts can bias analyses of low-abundance features; accordingly, we applied a conservative filter to remove genes with low expression (<75 reads in <4 samples). For bacteria and fungi, we applied a stringent e-value cutoff (<1e-6) in EggNOG to reduce false positives and misassigned plant reads (Supplemental Figure 2). These criteria yielded a substantial fraction of unknown reads, some attributable to protists and viruses, which were retained as unknown because they were outside the study’s scope. Unknown metatranscriptomic reads in this study and others, may harbor valuable information that could be revisited as annotation tools and databases improve.

### Rhizosphere microbial composition and comparison of sequencing approaches

Microbial community composition was calculated from three sources: active amplicon (16S from RNA), complete amplicon (16S from DNA), and transcriptomic sequencing. Beta-diversity analysis, conducted independently for each sequencing type, revealed highly similar patterns of microbiome composition driven by soil type and temperature treatment (Supplemental Figure 3). Each dataset was trimmed and collapsed at the genus level. Read counts for the transcriptomic dataset were up to 100-fold higher (∼15M) than for either the active amplicon (250K) or the complete amplicon (156K). For a fair comparison between sequencing types, samples were rarified to a minimum read count of 50K. After filtering and rarefaction, 356 genera were assigned in the active amplicon dataset, 336 in the complete amplicon dataset, and 913 bacterial genera in the transcriptome dataset (Supplemental Figure 4A). A shared set of 164 genera represented the majority of the community’s relative abundance in the complete amplicon (78%), the active amplicon (68%), and the transcriptomic data (65%).

Transcriptomic data also seemed to capture more variation due to host genotype than either amplicon dataset (Supplemental Figure 3D). When comparing shared genera at the mean value for each genotype-treatment level, we observed linear relationships between sequencing types. However, comparisons involving the transcriptomic dataset were right-skewed, indicating that low-abundance taxa are better captured by transcriptomic sequencing (Supplemental Figure 4B). Similarly, we observed lower variation between biological replicates in the transcriptomic dataset than in either amplicon dataset (Supplemental Figure 4C). The two amplicon datasets were most similar to each other, with slightly greater variability in the active 16S dataset than in the complete 16S dataset.

The lower variation in taxonomic assignment in the transcriptomic dataset may be partly due to its higher resolution compared to the amplicon dataset. Many genera identified in the transcriptomic dataset could not be distinguished by amplicon sequencing. For example, amplicon sequencing grouped *Paraburkholderia*, *Burkholderia*, and *Caballeronia,* annotated in the SILVA database as *Burkholderia-Caballeronia-Paraburkholderia,* whereas the transcriptomic dataset identified each genus separately. These genera are all members of the *Burkholderia sensu lato* group that share 96-100% sequence similarity of their published full-length 16S rRNA genes. Yet key functional differences exist among *Burkholderia sensu lato* members, including their roles in nitrogen and carbon cycling^66^.

Although we expected greater similarity between the active 16S and transcriptomic datasets, the two amplicon datasets were most similar to each other. This observation is likely driven by amplification and database bias, as both amplicon datasets were assigned taxonomy using the SILVA database, whereas the transcriptomics data were assigned using RefSeq. In almost all metrics (number of genera identified, taxonomic relative abundance, and replicate variance), the amplicon datasets performed similarly. However, among the 164 taxa that overlapped across the datasets, they accounted for a greater proportion of total abundance in the complete amplicon dataset than in the active amplicon and transcriptomics datasets (78% vs. 68% and 65%). This indicates that while the set of taxa identified by all three sequencing types dominates the community, activity and gene expression are more broadly distributed across both abundant and rare microbes, suggesting functional contributions are not proportional to microbial relative abundance.

The metatranscriptomic dataset provides both higher taxonomic resolution and functional insight, capturing the same community structure observed by amplicon sequencing while extending it to reveal which members are transcriptionally active and which pathways are enriched. Although more time- and resource-intensive, metatranscriptomics offers a comprehensive view of microbiome composition and function, making it an extremely informative approach for dissecting plant–microbe interactions.

### Maize and Sorghum Differential Expression Analysis

We compiled a joint maize–sorghum transcriptome dataset restricted to orthologous genes, with samples averaging 2.7 million reads assigned to orthologous genes. A complete list of orthologs used in this study is included in Supplemental Table 4. To confirm that combining datasets did not obscure the heat stress response, we compared our results with those from published maize and sorghum heat stress transcriptomic studies. Across six independent studies spanning multiple tissues and developmental stages, 67 genes were consistently differentially expressed in response to heat stress. Of these, 36 genes (53.7%) were also identified in our dataset (Figure 3A), including genes with demonstrated roles in heat tolerance: *hsp4*, *hsp7*, *hsp10*, and *ms42*^67,68^. Details of these studies, along with a complete list of identified genes, are available in Supplemental Datasets 1 and 2^69–73^.

**Figure 3.**
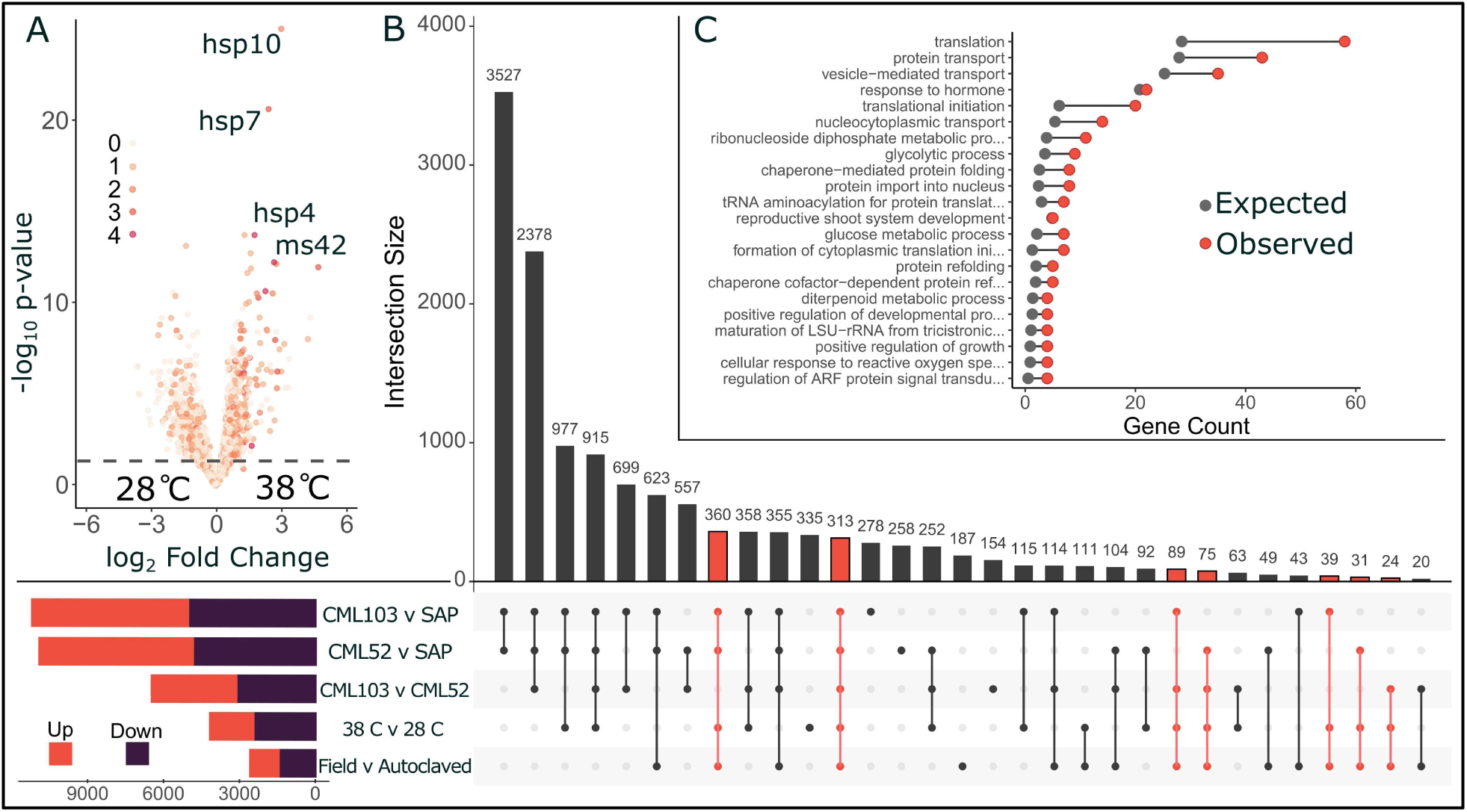
Shared and divergent heat stress responses in maize and sorghum. Differentially expressed genes under heat stress from a combined maize-sorghum ortholog dataset are displayed in a volcano plot, with points colored by the number of publicly available datasets in which they appear. Key heat-response genes, including multiple heat shock proteins (hsp4, hsp7, and hsp10) and a mitochondrial stress protein (ms42), are strongly upregulated and consistently recovered across datasets (A). A large number of differentially expressed genes were identified between the genotypes. To narrow our focus, we selected 931 genes that were differentially expressed (highlighted in red) when comparing temperatures, soil type, and genotype independently (B). GO term enrichment in the list of intersecting genes differentially expressed due to temperature, soil treatment, and genotype (C).

Differential expression analysis revealed extensive transcriptional divergence between genotypes, particularly between species (Figure 3B). To narrow the focus to genes involved in microbiome regulation in response to heat stress, we selected intersecting genes, those differentially expressed across genotype, temperature, and soil comparisons. Of the total list of ∼13,000 genes differentially expressed in at least one comparison, 931 intersected all three (Figure 3B highlighted in Red).

Functional enrichment analysis of these intersecting genes revealed a significant overrepresentation of translation- and transport-related processes, as well as chaperone-mediated protein folding and hormone responses (Figure 3C and Supplemental Dataset 1). These functions indicate a role of protein homeostasis and stress signaling in the microbe-mediated, shared maize-sorghum heat response. Our analysis used orthologous genes between maize and sorghum to efficiently integrate datasets. Noting that while syntenic orthologs are more likely to have conserved functions, including all orthologs allows us to capture genes with species-specific functional and expression differences that may be relevant under different soil conditions and high temperatures^74,75^.

### Microbial Gene and Pathway Enrichment

Microbial genes annotated by EggNOG showed global microbial gene expression was driven by soil type and temperature (Figure 4A). Though the overall community gene expression was dominated by these environmental parameters, host genotype was also a significant modulator of the microbial temperature response. 6,258 of 27,911 microbial genes were significantly differentially expressed in response to temperature across at least one genotype-by-soil grouping (Figure 4B). The expression patterns show some similarities between groups but reveal unique expression profiles driven by both soil type and genotype.

**Figure 4.**
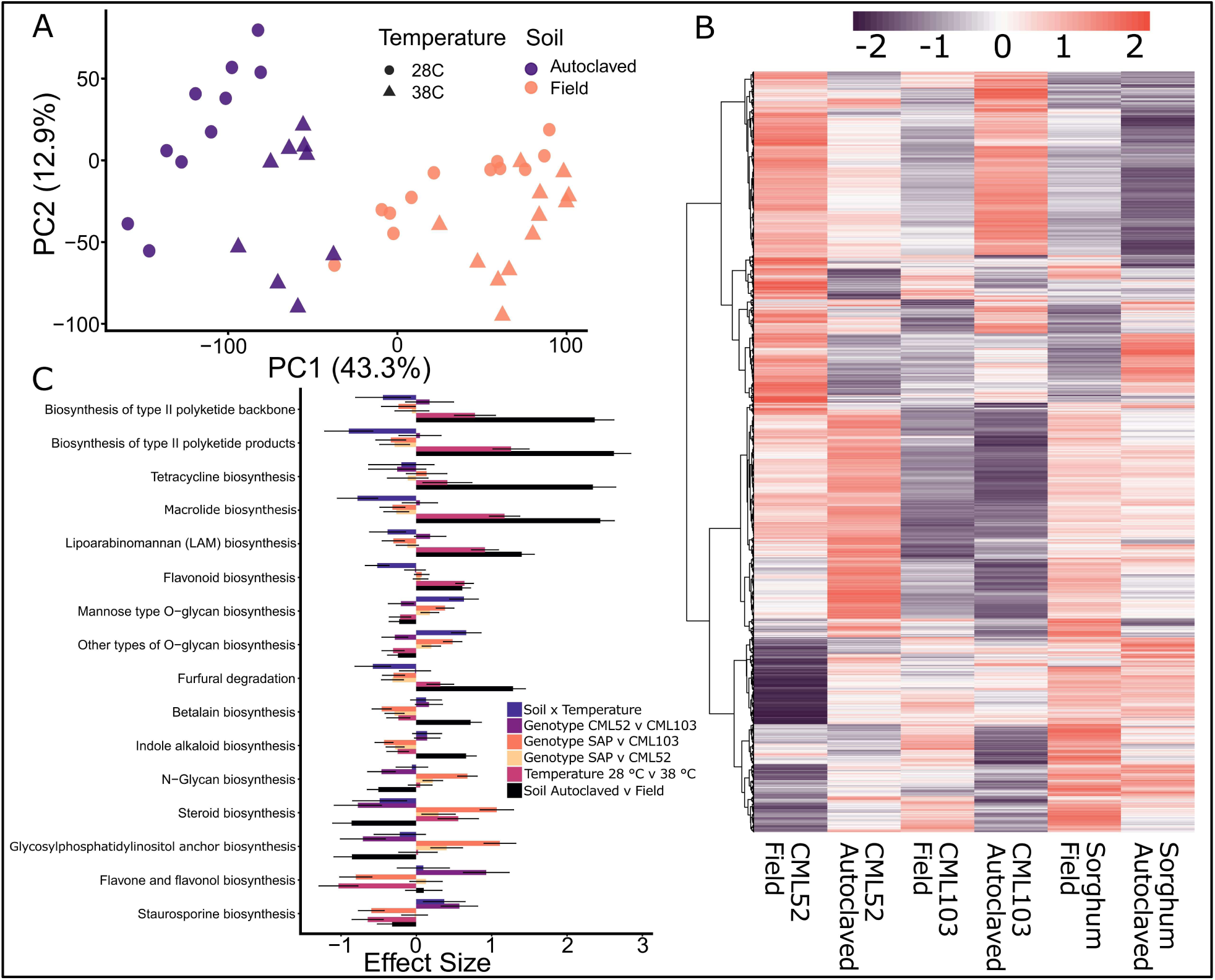
Microbial functional responses to soil treatment, temperature, and genotype. Principal component analysis of microbial gene expression across samples. Points are colored by soil treatment and shaped by temperature (circles = 28°C, triangles = 38°C)(A). Heatmap of normalized expression (Z-score) for differentially expressed (temperature) microbial genes across samples. Hierarchical clustering was performed on both pathways (rows) and samples (columns) using Euclidean distance. Red indicates higher relative expression, and purple indicates lower relative expression. Sample labels indicate genotype and soil type (B). Effect sizes of the top microbial pathways for main and interaction effects derived from ALDEx2 GLM. Bars represent estimated effect sizes for soil, temperature, genotype, and the interaction between soil and temperature. Error bars represent standard errors (C).

Microbial pathways were consistently differentially expressed in response to soil type and, in many cases, to temperature as well. Quantification of the main and interaction effects using an ALDEx2 GLM revealed that soil type was the dominant factor influencing the differential abundance of many metabolic pathways (Figure 4C). Specifically, the Soil contrast exhibited the largest effect sizes across several top pathways, including the biosynthesis of macrolides and polyketides, underscoring the overriding importance of the soil environment in shaping the microbial functional potential. Pathways such as furfural degradation and flavonoid biosynthesis showed strong effect sizes driven by the Soil x Temperature interaction, suggesting a condition-specific mechanism for the microbial degradation of aromatic and toxic compounds. The significant effect sizes observed for the biosynthesis of several antibiotic classes, including macrolides, tetracycline, and novobiocin, indicate that the immediate soil environment is the key trigger for the mobilization of microbial secondary metabolite gene clusters. These shifts in antimicrobial biosynthesis likely reflect a dynamic, environmentally driven competitive landscape crucial for the assembly and functional stability of the root-associated microbiome.

### Plant Gene and Microbial Pathway Coexpression

To quantify the linear association between the host and microbiome functional profiles, we performed a Mantel test on distance matrices of plant and microbial gene expression. This analysis revealed significant co-variation between host and microbe transcriptomes (Mantel r = 0.258, p = 2e-04; Figure 5A), demonstrating that changes in the microbiome transcriptome mirror shifts in plant gene expression. However, clustering of points in the Mantel test revealed relationships in the data driven in part by soil type and host species. To estimate the contribution of host species, temperature, and soil type to gene expression, we conducted a PERMANOVA analysis to assign numerical values to each variable for both plant and microbial gene expression. In both PERMANOA analyses, all main effects were statistically significant (p-value < 0.05). Soil type had a much greater effect on microbial gene expression, and host species had a greater effect on plant gene expression (Figure 5A).

**Figure 5.**
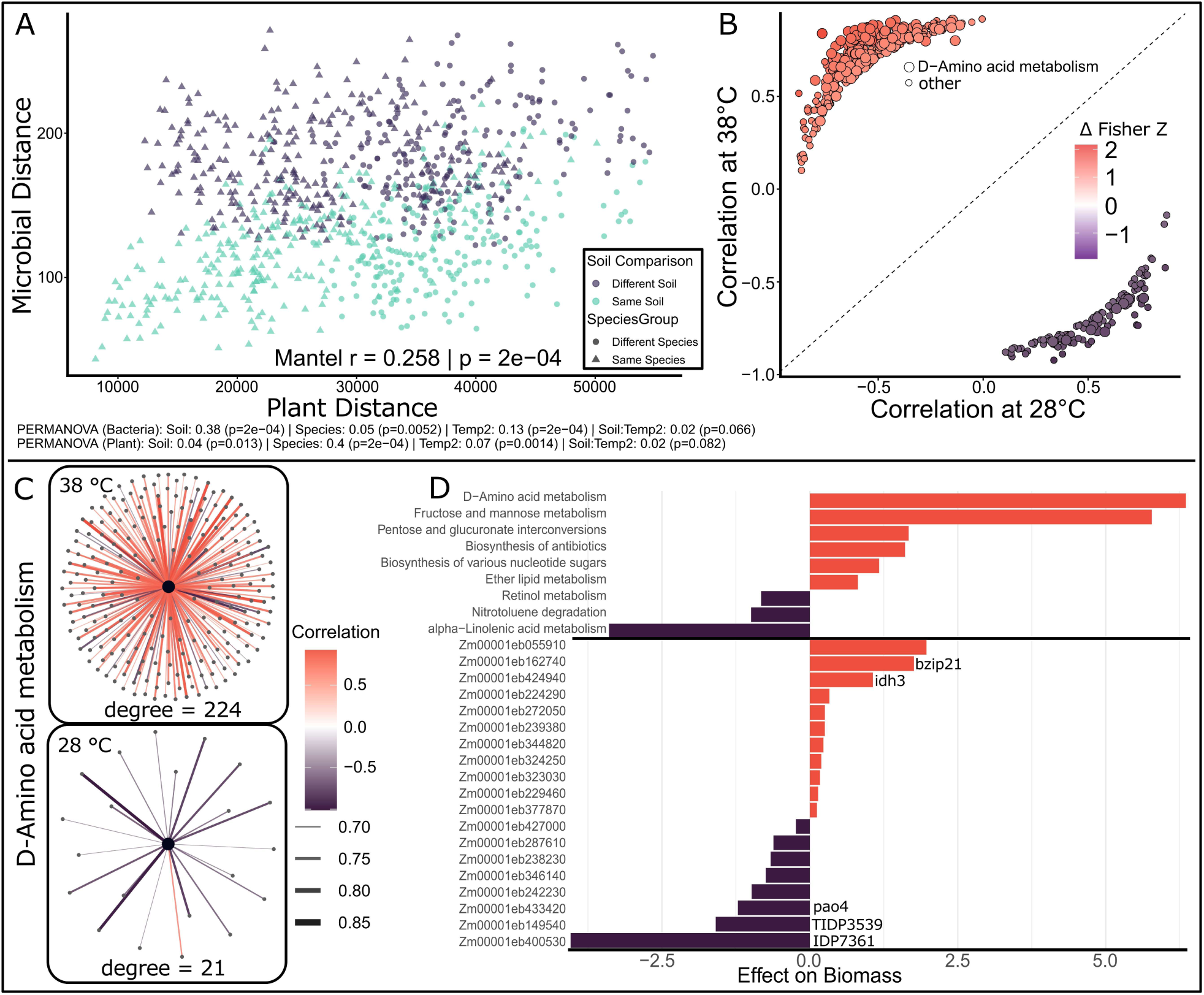
Co-expression between plant and microbial transcriptional profiles. Mantel test comparing pairwise distance matrices of plant and microbial gene expression, revealing significant co-variation between host and microbial transcriptomes. Clustering and PERMANOVA analysis indicate data structuring by soil type and host species (A). Differential bipartite correlation analysis showing the correlation values (Spearman’s Rho) for Microbial pathway-Plant gene pairs at high and optimal temperatures. Only points that were significantly differentially correlated (FDR < 0.05) are shown on the plot, colored by change in Z-score (B). Bipartite network showing D-amino acid metabolism pathway and connections (Spearman’s Rho > 0.7) to plant genes at optimal conditions and under heat stress (C). Elastic net regression identifying microbial pathways and plant genes most strongly associated with variation in biomass. Bars represent feature coefficients; positive values indicate increased expression associated with greater biomass (D).

To further explore specific co-regulated functions that responded to high temperatures, we constructed a bipartite co-expression network linking plant genes and microbial pathways at both optimal and high temperatures and conducted a differential correlation analysis to identify relationships between plant genes and microbial pathways that are different between temperatures (Δ Fisher Z ≥ 1 and FDR < 0.05; Figure 5B). The results of this analysis, shown in Figure 5B, demonstrate that most differentially correlated pairs fall in the upper-left quadrant, indicating that more plant–microbe relationships shifted from negative or weak associations under optimal conditions to strong positive correlations under heat stress (361 pairs) than shifted toward negative correlations (131 pairs). Under heat stress, clusters of genes and pathways that are often lowly expressed under optimal conditions are switched on. Of 492 correlated pathway and gene pairs, 245 contained the microbial D-amino acid metabolism pathway. Microbial D-amino acid metabolism exhibits a moderate degree of mostly negative correlations with plant genes under optimal conditions, whereas under heat stress, it shows a high degree of positive connectivity with maize gene expression (Figure 5C).

We employed elastic net regression to associate transcriptional features with phenotypic outcomes, identifying both plant genes and microbial pathways predictive of phenotypic variation in biomass, root volume, and heat stress tolerance (Figure 5D). The elastic net model identified 11 microbial pathways and 31 plant genes as explanatory of plant biomass, 8 microbial pathways and 39 plant genes for root volume, 2 microbial pathways and 49 plant genes for root shoot ratio, and 5 microbial pathways and 28 plant genes for heat stress.

Of the plant genes associated with heat tolerance biomarkers a putative palmitoyl protein thioesterase (*Zm00001eb055910*) emerged as the strongest positive predictor of biomass. As a regulator of protein depalmitoylation and membrane-associated signaling complexes, it plausibly influences root surface communication with the microbiome during heat stress. Two additional genes, *bZIP21* and *IDH3*, were also positively correlated with biomass. *bZIP21* integrates ROS, ABA, and carbon-allocation signals and may help maintain coordinated host–microbe signaling under stress, while *IDH3*, a mitochondrial NAD-dependent isocitrate dehydrogenase subunit, supports TCA-cycle activity, and organic acid production, functions consistent with enhanced metabolic capacity in heat-tolerant plants.

Conversely, *PAO4*, *TIDP3539*, and *IDP7361* were negatively associated with biomass. *PAO4* mediates polyamine catabolism, a pathway broadly activated under cellular stress; *TIDP3539* encodes an RNA-binding protein likely involved in post-transcriptional stress regulation; and *IDP7361* (mannose-1-phosphate guanyltransferase) contributes to GDP-mannose biosynthesis linked to cell-wall modification. Reduced expression of these stress-associated genes in high-biomass plants indicates a shift away from generalized stress signaling and toward metabolic efficiency and microbially supported growth.

Similar to the elastic net analysis, a random forest approach was applied to identify features that explain plant biomass, root volume, and heat stress. When both microbial pathways and plant genes were included, plant genes were rated higher variable importance than microbial pathways in all tests (Supplemental Figure 5). To identify a shortlist of candidate genes and pathways, the results of the three analyses were compiled, and features identified in two or more models were selected (Supplemental Figure 6 and Supplemental Table 5).

The same three analyses were conducted using microbial data at the gene level, with similar results, and generated a list of 73 candidate microbial genes and 81 plant genes identified by at least 2 tests (supplemental table 5). Notably, two microbial genes, *amt* from *Sphingobacteriales* and *gcvT* from *Planctomycetota*, were identified in four of the nine tests. Members of the order *Sphingobacteriales* are frequently implicated in plant stress responses^76^. Differential regulation of the *Sphingobacteriales amt* ammonium transporter suggests that nitrogen cycling may underlie *Sphingobacteriales*’ involvement in host stress biology. Similarly, the *gcvT* gene encodes a glycine cleavage T protein involved in C₁ metabolism and amino-acid turnover.

The candidate maize, sorghum ortholog supported by the most total analyses is *Zm00001eb373970*, named *arftf27* or *aux16*, an annotated auxin signalling transcription factor. This gene has previously been associated with environmental stress and root morphology, but not with microbiome modulation, although it is well known that auxin signalling is a primary mode of communication between plants and their root microbiomes^77,78^.

Several microbial pathways and maize gene relationships identified through co-expression analysis appear biologically plausible, particularly those linking microbial carbon and sugar metabolism (e.g., pentose and glucuronate interconversions, fructose and mannose metabolism, and starch and sucrose metabolism) with plant genes involved in the biosynthesis and transport of sugars and amino acids (Supplemental Figure 6). Such coupling is consistent with increased rhizodeposition under heat stress, in which elevated root exudation could promote the expression of carbon-catabolizing enzymes by microbes. Similarly, microbial retinol metabolism and terpenoid backbone biosynthesis were correlated with maize transcription factors associated with oxidative stress (e.g., ethylene-responsive and peroxidase genes), suggesting a shared role in modulating reactive oxygen balance or lipid-derived signaling during heat adaptation. The presence of microbial D-amino acid (DAA) metabolism, coupled with plant genes related to protein modification and turnover (e.g., serine/threonine kinases, SUMO-conjugating enzymes, and REF6 demethylase), may reflect active exchange or sensing of DAA in the rhizosphere. We focused on this pathway because, although DAA have been reported to exert both positive and negative effects on plants, the highly correlated expression profiles suggest a novel finding: that these molecules may participate directly in plant–microbe synergistic responses to abiotic stress^79–81^.

By integrating plant and microbial metatranscriptomes into a single framework, we can identify co-expression patterns of microbial and plant functions that were not apparent when each dataset was analyzed independently. Treating the plant and rhizosphere transcriptomes as interacting systems that equally contribute to the host phenotype has uncovered coordinated host–microbe responses to heat stress and provides a unique window into the functional coupling of plants and their rhizosphere microbiome under abiotic stress.

### Evidence for a Plant-Mediated DAA Metabolism Shift under Heat Stress

The observed correlation between microbial DAA metabolism and plant performance was genotype and soil-dependent (Supplemental figure 7A and 7B). The most consistent increase was in the heat-tolerant CML52, and the strongest increase was observed in CML103 individuals that displayed high biomass in the autoclaved soil (Figure 2A and Supplemental Figure 7B). This pattern suggests that the upregulation of microbial DAA metabolism is associated with plant performance under heat stress. The bulk soil control (no plant) showed no corresponding increase in DAA metabolism and trended slightly downward, though the limited sample size precluded statistical confirmation. Moreover, analysis of a publicly available metagenomic dataset from 30 grassland soils across Europe revealed a decrease in DAA metabolism under experimental heat stress (Supplemental Figure 7C)^82^. This is consistent with previous findings that DAA cycling tends to decline when plants and microbial communities are directly stressed^80^. Together, these results indicate that the observed increase in microbial DAA metabolism is unlikely to reflect a general stress response. Instead, it points toward a plant-mediated mechanism, in which root-derived metabolites alter the rhizosphere environment and selectively activate microbial DAA metabolic pathways in specific genotypes and soil contexts. Although our data strongly support this model, further mechanistic experimentation will be required to determine the causality and directionality of this interaction.

Maize and sorghum genes significantly correlated with microbial DAA metabolism support a signaling-based hypothesis rather than an independent stress-response or nutrient-transfer process (Supplemental Table 5). Notably, only 58 of the 224 plant genes correlated with DAA metabolism were also differentially expressed under heat stress. This is significantly fewer than expected if both responses were independently driven by temperature (one-tailed hypergeometric test, p < 0.01), suggesting that DAA metabolism and plant gene expression are functionally coupled. Highly correlated genes included key carbon metabolism enzymes such as invertase2 (*ivr2*), aldolase1 (*ald1*), enolase1 (*eno1*), and phosphoenolpyruvate carboxylase4 (*pep4*), consistent with increased metabolic flux through glycolysis and sucrose mobilization that could alter carbon exudation to stimulate rhizosphere colonization. Upregulation of redox-regulatory and calcium-signaling genes (*nucleoredoxin1*, *glutathione reductase2*, *prdx4*, *cipk20/33*, *annexin1/2*) suggests a ROS-activated signaling and transport network. The co-expression of transcription factors possibly linked to hormone and stress signaling (*WRKY*, *CAMTA*, and *ARF* families) could indicate integration of DAA-associated cues into regulatory pathways. Structural remodeling at the root–soil interface may also be involved, given correlations with cellulose synthase8 *(cesa8)*, xyloglucan endotransglucosylase/hydrolase7 *(xth7)*, and roothair defective3 *(rth3)*. Multiple of these gene types in maize and sorghum have previously been reported to affect microbiome function, especially the formation of root hairs by *rth3* and sugar allocation by *ivr2*^83–85^. Collectively, these expression patterns suggest that microbial DAA metabolism engages a maize signaling feedback loop involving carbon flux, redox homeostasis, and hormonal regulation, consistent with the hypothesis that DAA metabolism acts as a rhizosphere signal.

Upon re-running the tri-analysis (differential correlation, elastic net, and random forest) on only genes annotated to D-amino acid metabolism, we observed that many of these genes were lowly expressed (did not meet our original threshold of >250 reads in at least three samples). The top two candidate genes (identified in 4 of 9 tests) were a *S-adenosylmethionine decarboxylase* (*speH)* from *Burkholderiaceae* and a *Branched-chain-amino-acid aminotransferase (ilvE)* from *Streptosporangiales*. The identities of genes significant in at least 2 of 9 analyses point to functional patterns consistent with the broader DAA metabolic response, including several genes involved in D-amino acid synthesis and turnover, including amino acid racemases (*alr, asr, dhaa, murD/F/I)*, aspartate ammonia-lyase (*ansB*), D-amino acid dehydrogenase (*dadA)*, diaminopimelate epimerase (*dapF*), and peptidoglycan biosynthesis (*dapf*), spanning diverse lineages including *Burkholderiales*, *Comamonadaceae*, *Pseudomonas putida*, and *Rhizobiaceae*.

These genes link DAA cycling to cell wall remodeling, amino acid interconversion, and stress adaptation, suggesting that DAA metabolism could mediate shifts in microbial growth or signaling in addition to nutrient uptake. Because many of these genes were expressed near detection limits, pathway-level enrichment analysis provided the resolution to detect coordinated functional shifts that would have been missed by single-gene filtering. This highlights how collapsing expression data to pathway-level summaries can reveal global microbiome functional responses across complex microbial communities.

The parallel changes in maize gene expression and microbial DAA metabolism suggest a synergistic relationship rather than independent responses to the same environmental conditions. One possibility is that root exudation stimulates microbial DAA metabolism, which can generate reactive oxygen species that trigger plant redox and hormone signaling networks controlling carbon distribution and root growth. Alternatively, both partners may be responding to a shared rhizosphere signal, such as local shifts in pH or redox potential driven by root exudation. While the coordinated expression patterns observed here support the existence of causal plant–microbe interactions, determining the direction and mechanisms of these exchanges will require future experimentation.

## Conclusion

Metatranscriptomics is a powerful alternative for microbiome profiling, revealing both the community’s composition and its active functional potential, while capturing community structure similarly to amplicon sequencing. Transcriptomics also enables us to treat the host and microbial transcriptome as a single system; our approach uncovers cross-kingdom interactions that connect microbial activity with plant responses to environmental stress. The systems-level, integrated host-microbe metatranscriptomic approach can be applied to many other host-microbiome systems to study the molecular basis of coordinated responses to the environment.

Across all analyses, soil treatment (autoclaved versus field soil) was the main influence on microbiome composition and function. While this method provided a clear experimental contrast, it may have been an overly severe disruption of native microbial communities. Nevertheless, the effects of temperature, genotype, soil treatment, and their interactions could be observed in the rhizosphere microbiome.

Our novel integrated host-microbe metatranscriptomic analyses revealed consistent effects of temperature, genotype, and their interactions on both the plant and microbial transcriptome. We identified a core set of plant genes—conserved between maize and sorghum—that respond to heat stress in tandem with microbial functional pathways. These candidate genes are promising targets for improving plant–microbe communication and enhancing thermotolerance through breeding or biotechnology. The pathway-level analysis of microbial transcriptomic profiles allowed us to resolve community-level mechanisms, capturing microbiome functions that individual genes alone could not reveal.

The coordinated upregulation of microbial D-amino acid (DAA) metabolism with maize genes involved in signaling, redox balance, and carbohydrate flux suggests a novel plant-mediated synergistic feedback loop critical for genotype-specific heat adaptation. We propose that DAA turnover may act as a rhizosphere signaling cue, warranting future mechanistic experiments to determine the direction and causality of this molecular exchange. While DAA metabolism emerged as a key node, correlated shifts in microbial carbon and redox metabolism with plant carbohydrate and signaling genes indicate that multiple mechanisms contribute to plant/rhizosphere adaptation under heat stress.

As other studies have shown, abiotic stress causes plants to divert resources to recruit beneficial microbes; our results add to the growing body of evidence that rhizosphere microbiome regulation is an important response to abiotic stress and are among the few to generate mechanistic hypotheses.^14,86,87^ To our knowledge, this is among the first studies to integrate plant and microbiome metatranscriptomes as a unified system under abiotic stress, revealing emergent patterns of cross-kingdom coordination. Collectively, these results support the Genotype × Environment × Rhizosphere Microbiome (GERMs) framework, demonstrating that plant adaptation results from the interplay of host genetics, environmental stimuli, and microbial function. By integrating host and microbial transcriptomes, this study provides a foundation for identifying breeding targets to develop next-generation crops capable of regulating their microbiomes to withstand a changing climate.

## Supporting information

Descriptions of Supplementals

## Acknowledgements

This work was supported by the National Institute of General Medical Sciences of the National Institutes of Health under award number R35GM151048, USDA NIFA Hatch award number 7002327, and seed funding from the NCSU Genetics and Genomics Academy. N. Korth was supported by a USDA-NIFA postdoctoral fellowship 2025-67012-44807. We thank Amy Grunden for providing field soil, Christine Hawkes for providing guidance and materials for the project, the NCSU Phytotron Staff for assistance with growth chambers, and the North Carolina State University High Performance Computing Services Core Facility (RRID:SCR_022168). Portions of the text were refined for clarity and style using OpenAI’s GPT-5 language model. The authors reviewed and verified all AI-assisted content to ensure accuracy and integrity of the scientific conclusions.

## Data availability

Sequenced transcriptomic data are available as raw fastq files deposited in the NCBI SRA database (project accession PRJNA1376519). A detailed bioinformatic pipeline used in this study is available at https://github.com/NateKorth/MicrobeRNAseq

## Notes

### Competing Interest Statement

The authors have declared no competing interest.

## References

1. R. E. ComStock & R. H. Moll. Genotype x Environment Interactions. Statistical genetics and plant breeding 982, 164–196 (1963).

2. Steiner, F., Zuffo, A. M., Teodoro, P. E., Aguilera, J. G. & Teodoro, L. P. R. Multivariate adaptability and stability of soya bean genotypes for abiotic stresses. Journal of Agronomy and Crop Science 207, 354–361 (2021).

3. Vandenkoornhuyse, P., Quaiser, A., Duhamel, M., Le Van, A. & Dufresne, A. The importance of the microbiome of the plant holobiont. New Phytologist 206, 1196–1206 (2015).

4. Addison, S. L., Rúa, M. A., Smaill, S. J., Singh, B. K. & Wakelin, S. A. Partner or perish: tree microbiomes and climate change. Trends in Plant Science 29, 1029–1040 (2024).

5. Plett, K. L., Bithell, S. L., Dando, A. & Plett, J. M. Chickpea shows genotype-specific nodulation responses across soil nitrogen environment and root disease resistance categories. BMC Plant Biology 21, 310 (2021).

6. Vaidya, P. & Stinchcombe, J. R. The Potential for Genotype-by-Environment Interactions to Maintain Genetic Variation in a Model Legume–Rhizobia Mutualism. Plant Comm 1, (2020).

7. Oyserman, B. O. et al. Extracting the GEMs: Genotype, Environment, and Microbiome Interactions Shaping Host Phenotypes. Front. Microbiol. 11, (2021).

8. Hultgren, A. et al. Impacts of climate change on global agriculture accounting for adaptation. Nature 642, 644–652 (2025).

9. Lindsey, R., Dahlman, L. & Blunden, R. J. Climate Change: Global Temperature. Climate Change.

10. Delouche, J. C. & Baskin, C. C. Accelerated Aging Techniques for Predicting the Relative Storability of Seed Lots.

11. Maitra, S. et al. Thermotolerant Soil Microbes and Their Role in Mitigation of Heat Stress in Plants. in Soil Microbiomes for Sustainable Agriculture: Functional Annotation (ed. Yadav, A. N.) 203–242 (Springer International Publishing, Cham, 2021). doi:10.1007/978-3-030-73507-4_8.

12. Abd El-Daim, I. A., Bejai, S. & Meijer, J. Improved heat stress tolerance of wheat seedlings by bacterial seed treatment. Plant Soil 379, 337–350 (2014).

13. Chai, Y. N. & Schachtman, D. P. Root exudates impact plant performance under abiotic stress. Trends in Plant Science 27, 80–91 (2022).

14. Tiziani, R. et al. Drought, heat, and their combination impact the root exudation patterns and rhizosphere microbiome in maize roots. Environmental and Experimental Botany 203, 105071 (2022).

15. Awika, J. M. Major Cereal Grains Production and Use around the World. in Advances in Cereal Science: Implications to Food Processing and Health Promotion vol. 1089 1–13 (American Chemical Society, 2011).

16. Food and Agriculture Organization of the United Nation. FAOSTAT statistical database. (2020).

17. Swigoňová, Z. et al. Close Split of Sorghum and Maize Genome Progenitors. Genome Res. 14, 1916–1923 (2004).

18. Chopra, R., Burow, G., Burke, J. J., Gladman, N. & Xin, Z. Genome-wide association analysis of seedling traits in diverse Sorghum germplasm under thermal stress. BMC Plant Biol 17, 12 (2017).

19. Abreha, K. B. et al. Sorghum in dryland: morphological, physiological, and molecular responses of sorghum under drought stress. Planta 255, 20 (2021).

20. Hao, L. et al. Genome-wide identification and comparative analysis of drought related genes in roots of two maize inbred lines with contrasting drought tolerance by RNA sequencing. Journal of Integrative Agriculture 19, 449–464 (2020).

21. Brisson, V. L. et al. Phosphate Availability Modulates Root Exudate Composition and Rhizosphere Microbial Community in a Teosinte and a Modern Maize Cultivar. Phytobiomes Journal 6, 83–94 (2022).

22. Huttenhower, C. et al. Structure, function and diversity of the healthy human microbiome. Nature 486, 207–214 (2012).

23. Walters, W. A. et al. Large-scale replicated field study of maize rhizosphere identifies heritable microbes. Proceedings of the National Academy of Sciences 115, 7368–7373 (2018).

24. Schultz, C. R., Desai, H. & Wallace, J. G. The Landscape of Maize-Associated Bacteria and Fungi Across the United States. 2023.07.11.548569 Preprint at 10.1101/2023.07.11.548569 (2023).

25. Gao, J. et al. Linkage mapping and genome-wide association reveal candidate genes conferring thermotolerance of seed-set in maize. Journal of Experimental Botany 70, 4849–4864 (2019).

26. McNellie, J. P., Chen, J., Li, X. & Yu, J. Genetic Mapping of Foliar and Tassel Heat Stress Tolerance in Maize. Crop Science 58, 2484–2493 (2018).

27. Zhang, M.-Y. et al. Relatively Low Light Intensity Promotes Phosphorus Absorption and Enhances the Ethylene Signaling Component EIN3 in Maize, Wheat, and Oilseed Rape. Agronomy 12, 427 (2022).

28. Xiong, C. et al. Plant developmental stage drives the differentiation in ecological role of the maize microbiome. Microbiome 9, 171 (2021).

29. Chen, L. et al. Designing specific bacterial 16S primers to sequence and quantitate plant endo-bacteriome. Sci China Life Sci 65, 1000–1013 (2022).

30. Apprill, A., McNally, S., Parsons, R. & Weber, L. Minor revision to V4 region SSU rRNA 806R gene primer greatly increases detection of SAR11 bacterioplankton. Aquat. Microb. Ecol. 75, 129–137 (2015).

31. Glöckner, F. O. et al. 25 years of serving the community with ribosomal RNA gene reference databases and tools. Journal of Biotechnology 261, 169–176 (2017).

32. Quast, C. et al. The SILVA ribosomal RNA gene database project: improved data processing and web-based tools. Nucleic Acids Research 41, D590–D596 (2013).

33. McMurdie, P. J. & Holmes, S. phyloseq: An R Package for Reproducible Interactive Analysis and Graphics of Microbiome Census Data. PLOS ONE 8, e61217 (2013).

34. Jari Oksanen et al. vegan: Community Ecology Package. (2025).

35. Bolger, A. M., Lohse, M. & Usadel, B. Trimmomatic: a flexible trimmer for Illumina sequence data. Bioinformatics 30, 2114–2120 (2014).

36. Morales, J. et al. A joint NCBI and EMBL-EBI transcript set for clinical genomics and research. Nature 604, 310–315 (2022).

37. Kim, D., Paggi, J. M., Park, C., Bennett, C. & Salzberg, S. L. Graph-based genome alignment and genotyping with HISAT2 and HISAT-genotype. Nat Biotechnol 37, 907–915 (2019).

38. Deng, Z.-L., Münch, P. C., Mreches, R. & McHardy, A. C. Rapid and accurate identification of ribosomal RNA sequences via deep learning. Nucleic Acids Research 50, e60 (2022).

39. Hufford, M. B. et al. De novo assembly, annotation, and comparative analysis of 26 diverse maize genomes. Science 373, 655–662 (2021).

40. Bo Wang et al. Pan-genome Analysis in Sorghum Highlights the Extent of Genomic Variation and Sugarcane Aphid Resistance Genes. Cold Spring Harbor Laboratory 2021.

41. Liao, Y., Smyth, G. K. & Shi, W. featureCounts: an efficient general purpose program for assigning sequence reads to genomic features. Bioinformatics 30, 923–930 (2014).

42. Emms, D. M. & Kelly, S. OrthoFinder: phylogenetic orthology inference for comparative genomics. Genome Biology 20, 238 (2019).

43. R Core Team. R: A language and environment for statistical computing. (2021).

44. Love, M. I., Huber, W. & Anders, S. Moderated estimation of fold change and dispersion for RNA-seq data with DESeq2. Genome Biology 15, 550 (2014).

45. Yates, A. D. et al. Ensembl Genomes 2022: an expanding genome resource for non-vertebrates. Nucleic Acids Res 50, D996–D1003 (2022).

46. Adrian Alexa & Jorg Rahnenfuhrer. topGO. Bioconductor http://bioconductor.org/packages/topGO/ (2025).

47. Kinsella, R. J. et al. Ensembl BioMarts: a hub for data retrieval across taxonomic space. Database (Oxford) 2011, bar030 (2011).

48. Wood, D. E., Lu, J. & Langmead, B. Improved metagenomic analysis with Kraken 2. Genome Biology 20, 257 (2019).

49. O’Leary, N. A. et al. Reference sequence (RefSeq) database at NCBI: current status, taxonomic expansion, and functional annotation. Nucleic Acids Res 44, D733–745 (2016).

50. Huerta-Cepas, J. et al. eggNOG 5.0: a hierarchical, functionally and phylogenetically annotated orthology resource based on 5090 organisms and 2502 viruses. Nucleic Acids Research 47, D309–D314 (2019).

51. Kanehisa, M., Furumichi, M., Sato, Y., Matsuura, Y. & Ishiguro-Watanabe, M. KEGG: biological systems database as a model of the real world. Nucleic Acids Res 53, D672–D677 (2025).

52. Buchfink, B., Reuter, K. & Drost, H.-G. Sensitive protein alignments at tree-of-life scale using DIAMOND. Nat Methods 18, 366–368 (2021).

53. Tenenbaum, D. & Maintainer, B. KEGGREST: Client-side REST access to the Kyoto Encyclopedia of Genes and Genomes (KEGG). Bioconductor http://bioconductor.org/packages/KEGGREST/ (2024).

54. Wickham, H. et al. Welcome to the Tidyverse. Journal of Open Source Software 4, 1686 (2019).

55. Boogaart, K. G. van den, Tolosana-Delgado, R. & Bren, M. compositions: Compositional Data Analysis. (2025).

56. Fernandes, A. D., Macklaim, J. M., Linn, T. G., Reid, G. & Gloor, G. B. ANOVA-Like Differential Expression (ALDEx) Analysis for Mixed Population RNA-Seq. PLOS ONE 8, e67019 (2013).

57. Gloor, G. B., Macklaim, J. M. & Fernandes, A. D. Displaying Variation in Large Datasets: Plotting a Visual Summary of Effect Sizes. Journal of Computational and Graphical Statistics 25, 971–979 (2016).

58. Tay, J. K., Narasimhan, B. & Hastie, T. Elastic Net Regularization Paths for All Generalized Linear Models. Journal of Statistical Software 106, 1–31 (2023).

59. Leo Breiman, Adele Cutler, Andy Liaw, & Matthew Wiener. randomForest: Breiman and Cutlers Random Forests for Classification and Regression. (2024).

60. Kassambara, A. ggpubr: ‘ggplot2’ Based Publication Ready Plots. (2023).

61. Wickham, H. ggplot2. WIREs Computational Statistics 3, 180–185 (2011).

62. Calleja-Cabrera, J., Boter, M., Oñate-Sánchez, L. & Pernas, M. Root Growth Adaptation to Climate Change in Crops. Front. Plant Sci. 11, (2020).

63. Diener, C. et al. Non-responder phenotype reveals apparent microbiome-wide antibiotic tolerance in the murine gut. Commun Biol 4, 316 (2021).

64. Olimi, E. et al. Plant microbiome responses to bioinoculants and volatiles. Environmental Microbiome 20, 55 (2025).

65. Li, H.-R. et al. Differential responses of root and leaf-associated microbiota to continuous monocultures. Environ Microbiome 20, 13 (2025).

66. Bach, E., Volpiano, C. G., Sant’Anna, F. H. & Passaglia, L. M. P. Genome-based taxonomy of Burkholderia sensu lato: Distinguishing closely related species. Genet Mol Biol 46, e20230122.

67. Diogo-Jr., R., et al. Maize heat shock proteins—prospection, validation, categorization and in silico analysis of the different ZmHSP families. Stress Biol 3, 37 (2023).

68. Li, Y. et al. Heat shock protein 101 contributes to the thermotolerance of male meiosis in maize. Plant Cell 34, 3702–3717 (2022).

69. Li, Y. et al. Transcriptomic Analysis Revealed the Common and Divergent Responses of Maize Seedling Leaves to Cold and Heat Stresses. Genes (Basel) 11, 881 (2020).

70. Yang, X. et al. Primary root response to combined drought and heat stress is regulated via salicylic acid metabolism in maize. BMC Plant Biology 22, 417 (2022).

71. Wang, T. et al. Single-cell transcriptomes reveal spatiotemporal heat stress response in maize roots. Nat Commun 16, 177 (2025).

72. Johnson, S. M. et al. Transcriptomic analysis of Sorghum bicolor responding to combined heat and drought stress. BMC Genomics 15, 456 (2014).

73. Ali, A. E. E. et al. Proteomic dataset of sorghum leaf and root responses to single and combined drought and heat stress. Sci Data 12, 97 (2025).

74. Zhang, Y. et al. Differentially Regulated Orthologs in Sorghum and the Subgenomes of Maize. Plant Cell 29, 1938–1951 (2017).

75. Adhikari, P., Mideros, S. X. & Jamann, T. M. Differential Regulation of Maize and Sorghum Orthologs in Response to the Fungal Pathogen Exserohilum turcicum. Front. Plant Sci. 12, (2021).

76. Hagaggi, N. Sh. A. & Abdul-Raouf, U. M. Drought-tolerant Sphingobacterium changzhouense Alv associated with Aloe vera mediates drought tolerance in maize (Zea mays). World J Microbiol Biotechnol 38, 248 (2022).

77. Zhang Kun et al. Identification of root morphology and discovery of candidate genes in maize inbred lines at the seedling stage. Journal of Plant Genetic Resources https://chn.oversea.cnki.net/kcms/detail/detail.aspx?filename=ZWYC202401008&dbcode=CJFQ&dbname=CJFDLAST2024&uniplatform=NZKPT (2024).

78. Saidi, A. & Hajibarat, Z. Computational study of environmental stress-related transcription factor binding sites in the promoter regions of maize auxin response factor (ARF) gene family. Notulae Scientia Biologicae 12, 646–657 (2020).

79. Gu, S.-X., Wang, H.-F., Zhu, Y.-Y. & Chen, F.-E. Natural Occurrence, Biological Functions, and Analysis of D-Amino Acids. Pharmaceutical Fronts 02, e79–e87 (2020).

80. Porras-Dominguez, J., Lothier, J., Limami, A. M. & Tcherkez, G. d-amino acids metabolism reflects the evolutionary origin of higher plants and their adaptation to the environment. Plant, Cell & Environment 47, 1503–1512 (2024).

81. Hill, P. W. et al. Acquisition and Assimilation of Nitrogen as Peptide-Bound and D-Enantiomers of Amino Acids by Wheat. PLOS ONE 6, e19220 (2011).

82. Knight, C. G. et al. Soil microbiomes show consistent and predictable responses to extreme events. Nature 636, 690–696 (2024).

83. Hartwig, R. P. et al. Drought response of the maize plant–soil–microbiome system is influenced by plant size and presence of root hairs. Ann Bot mcaf033 (2025) doi:10.1093/aob/mcaf033.

84. Korth, N. et al. Genomic co-localization of variation affecting agronomic and human gut microbiome traits in a meta-analysis of diverse sorghum. G3: Genes, Genomes, Genetics jkae145 (2024).

85. Sun, K. et al. Nitrogen fertilizer-regulated plant-fungi interaction is related to root invertase-induced hexose generation. FEMS Microbiol Ecol 96, fiaa139 (2020).

86. Liu, J., Zhen, B., Qiu, H., Zhou, X. & Zhang, H. Impact of waterlogging and heat stress on rice rhizosphere microbiome assembly and potential function in carbon and nitrogen transformation. Archives of Agronomy and Soil Science 69, 1920–1932 (2023).

87. Vescio, R., Malacrinò, A., Bennett, A. E. & Sorgonà, A. Single and combined abiotic stressors affect maize rhizosphere bacterial microbiota. Rhizosphere 17, 100318 (2021).

