## Supplementary figures and images for "Investigating GERMs: How Genotype, Environment, and Rhizosphere Microbiome interactions underlie heat response in maize and sorghum"

### Supplemental Figure 1

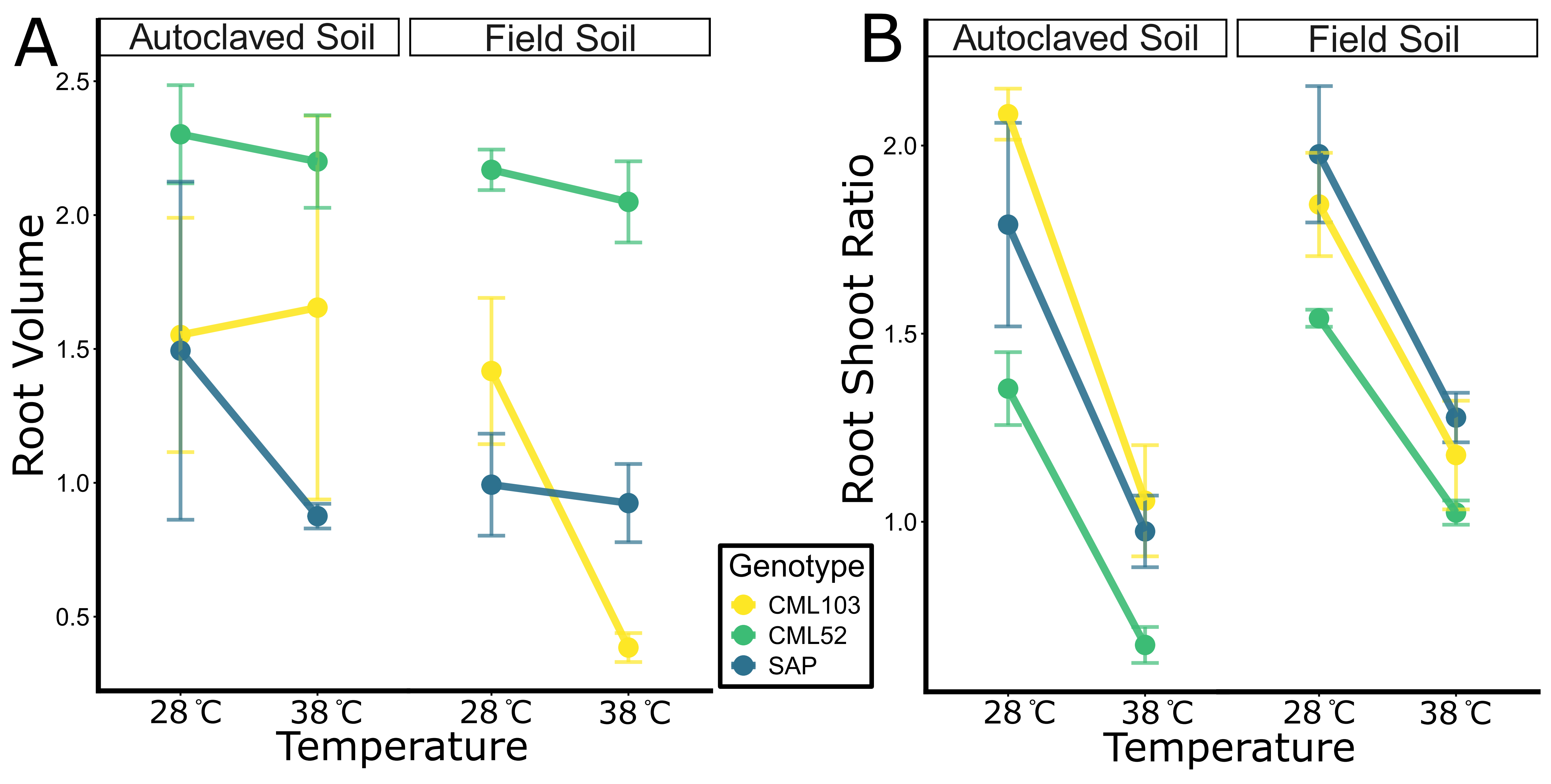

### Supplemental Figure 2

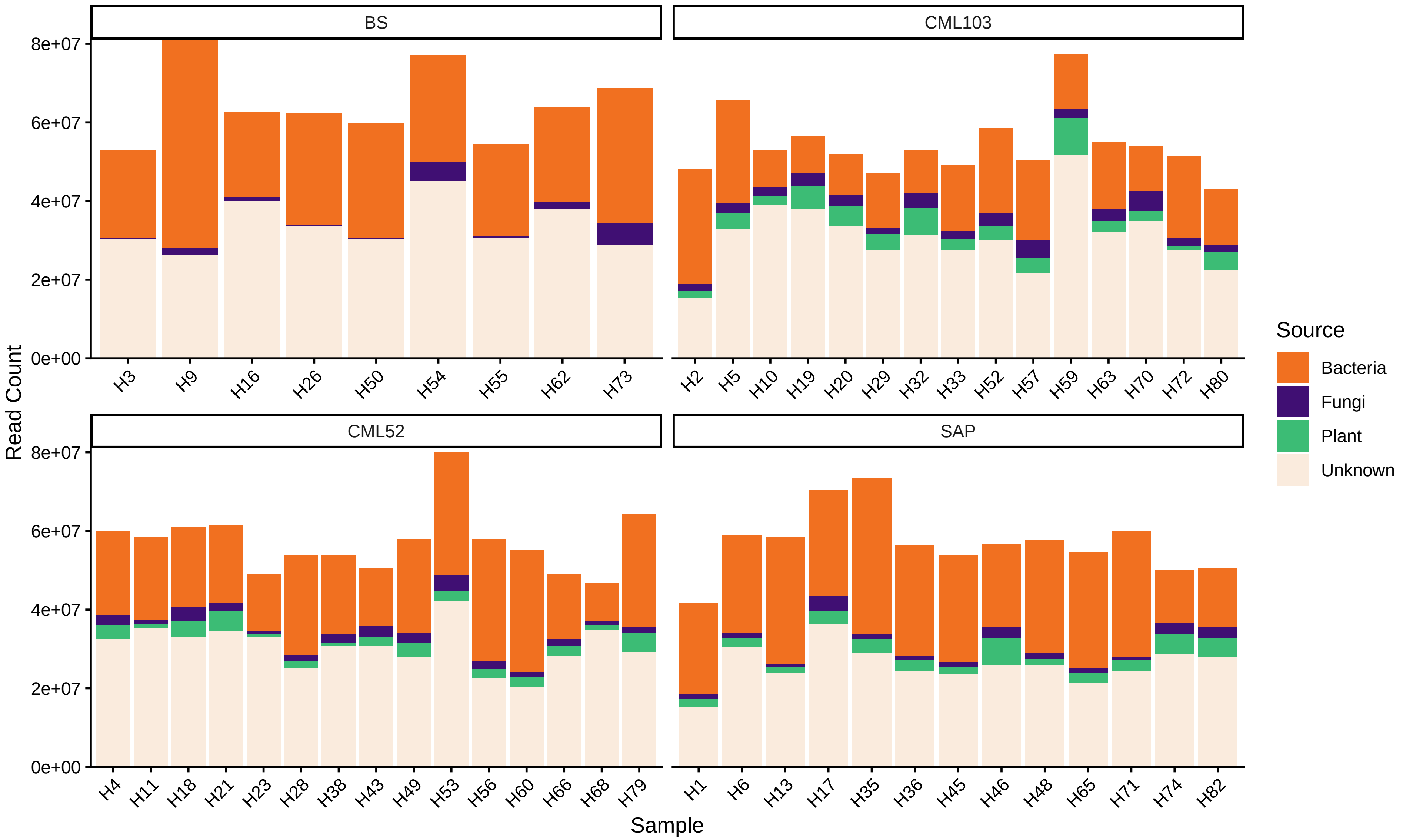

### Supplemental Figure 3

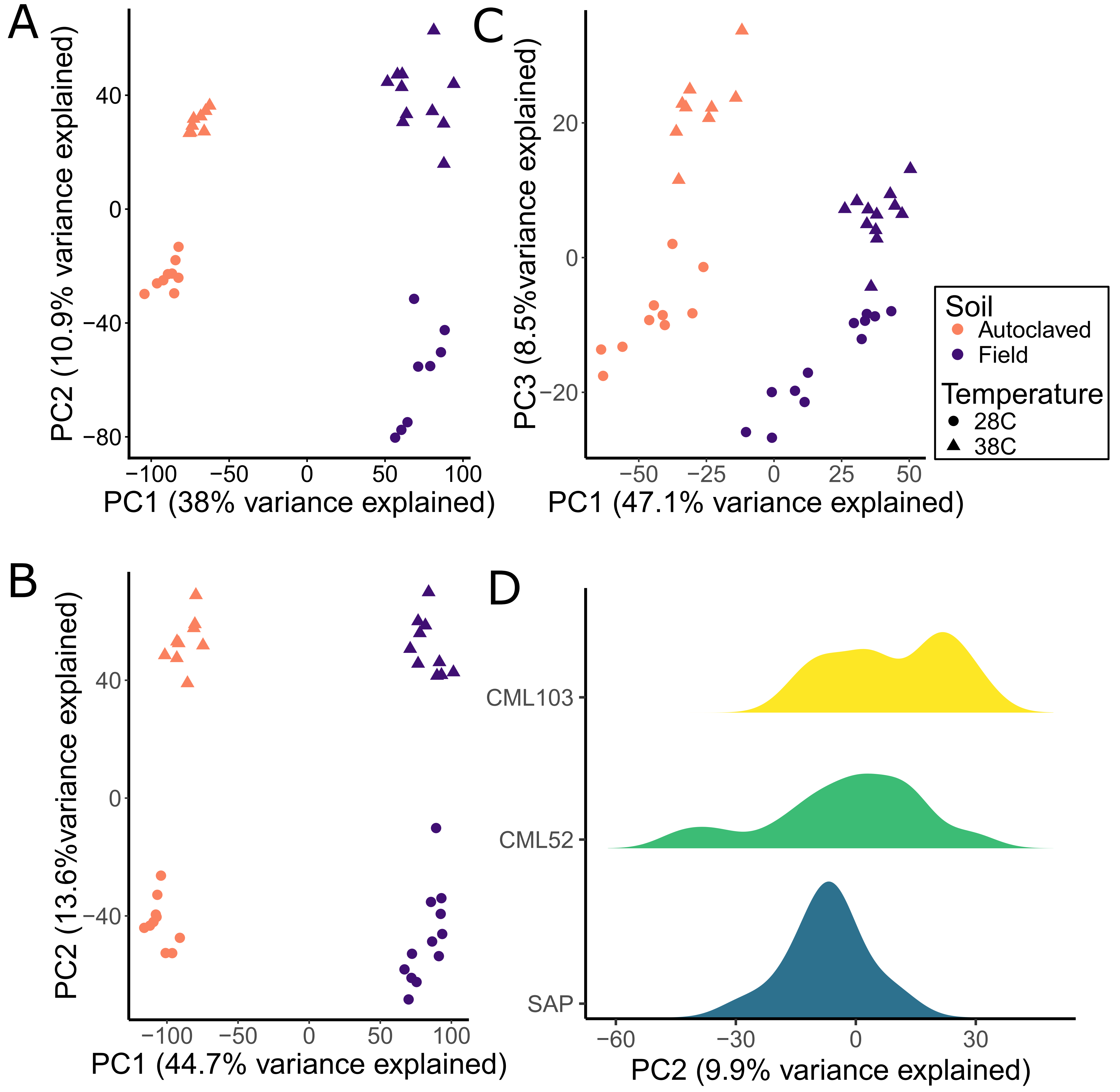

### Supplemental Figure 4

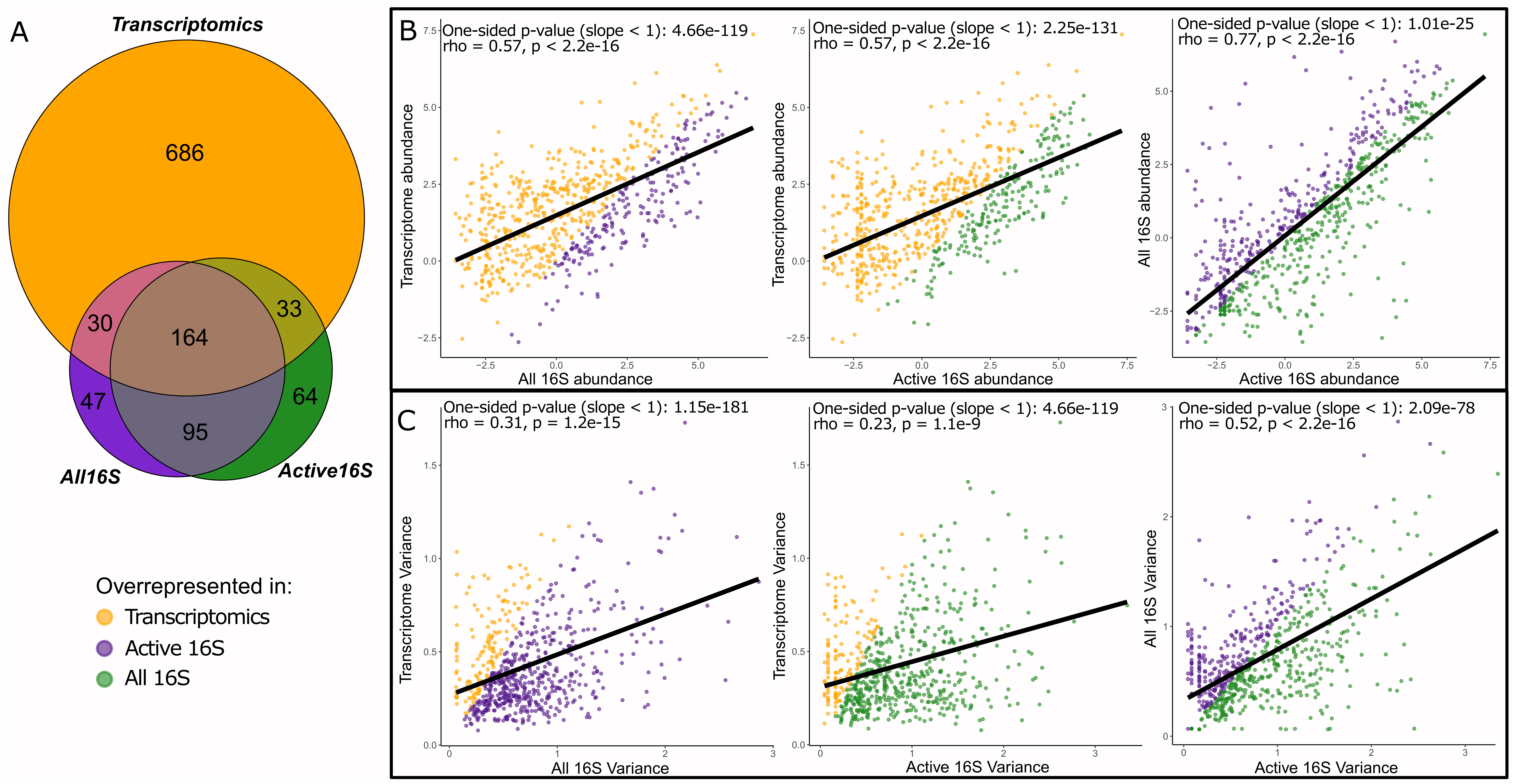

### Supplemental Figure 5

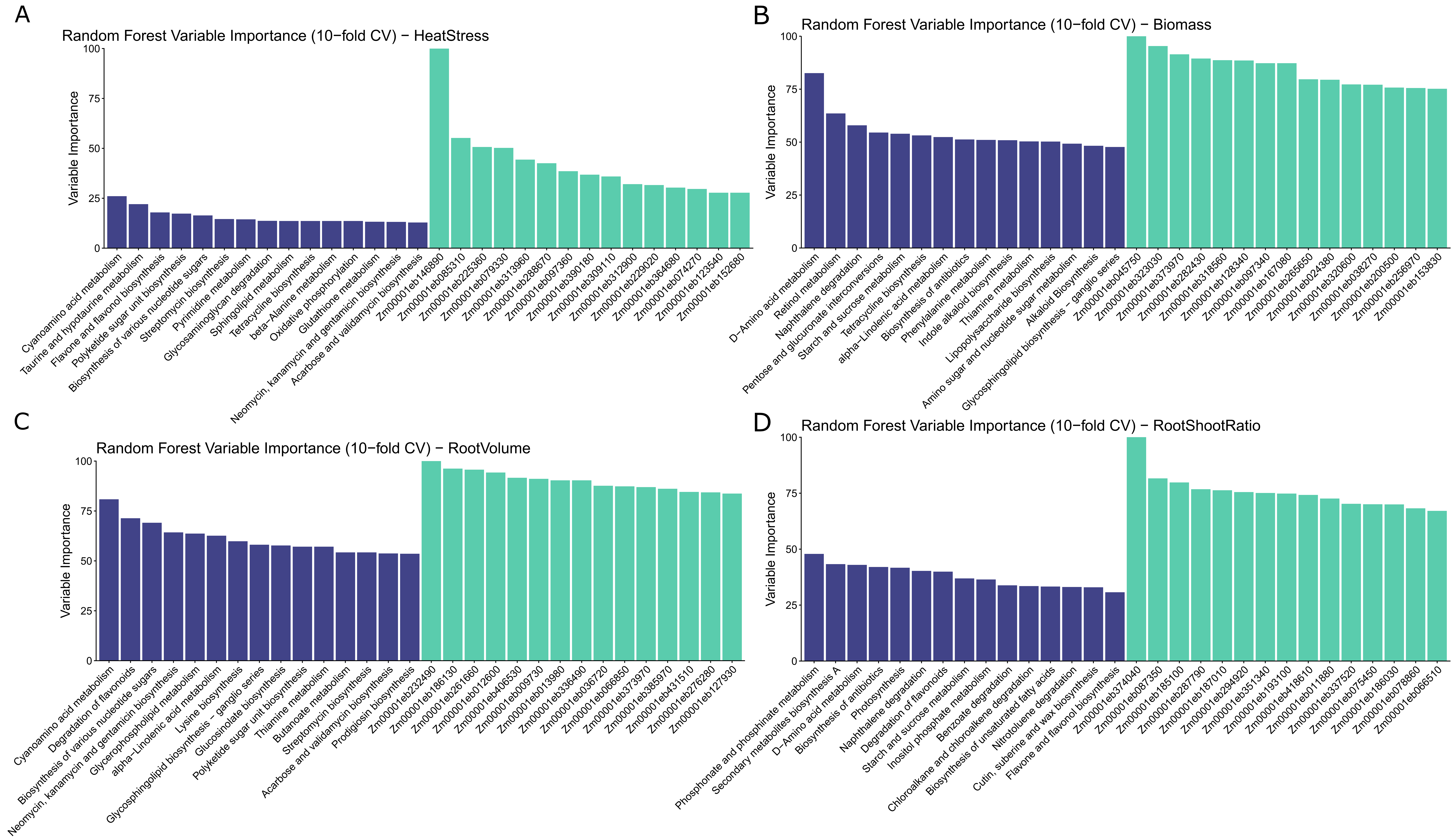

### Supplemental Figure 6

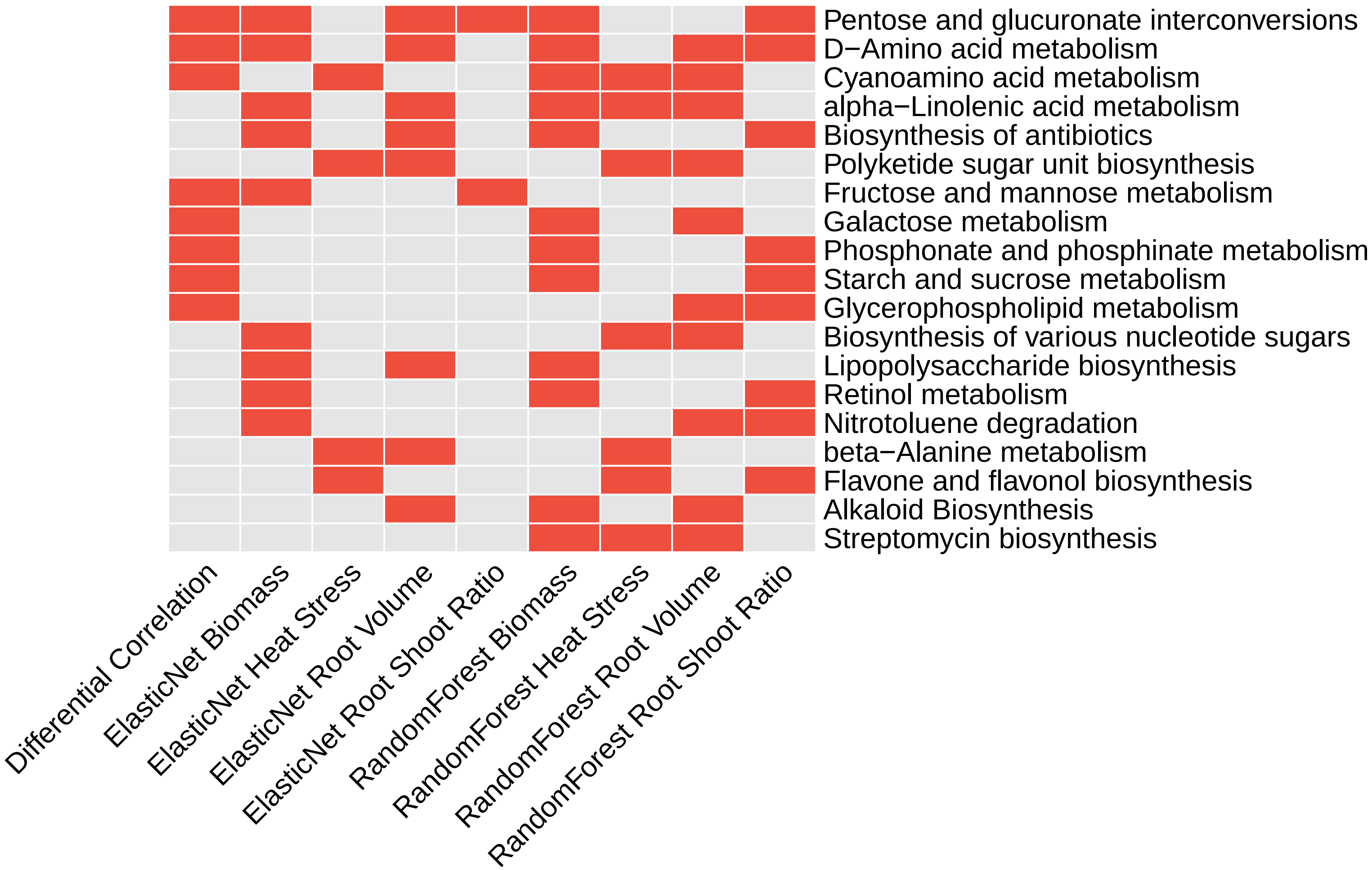

### Supplemental Figure 7

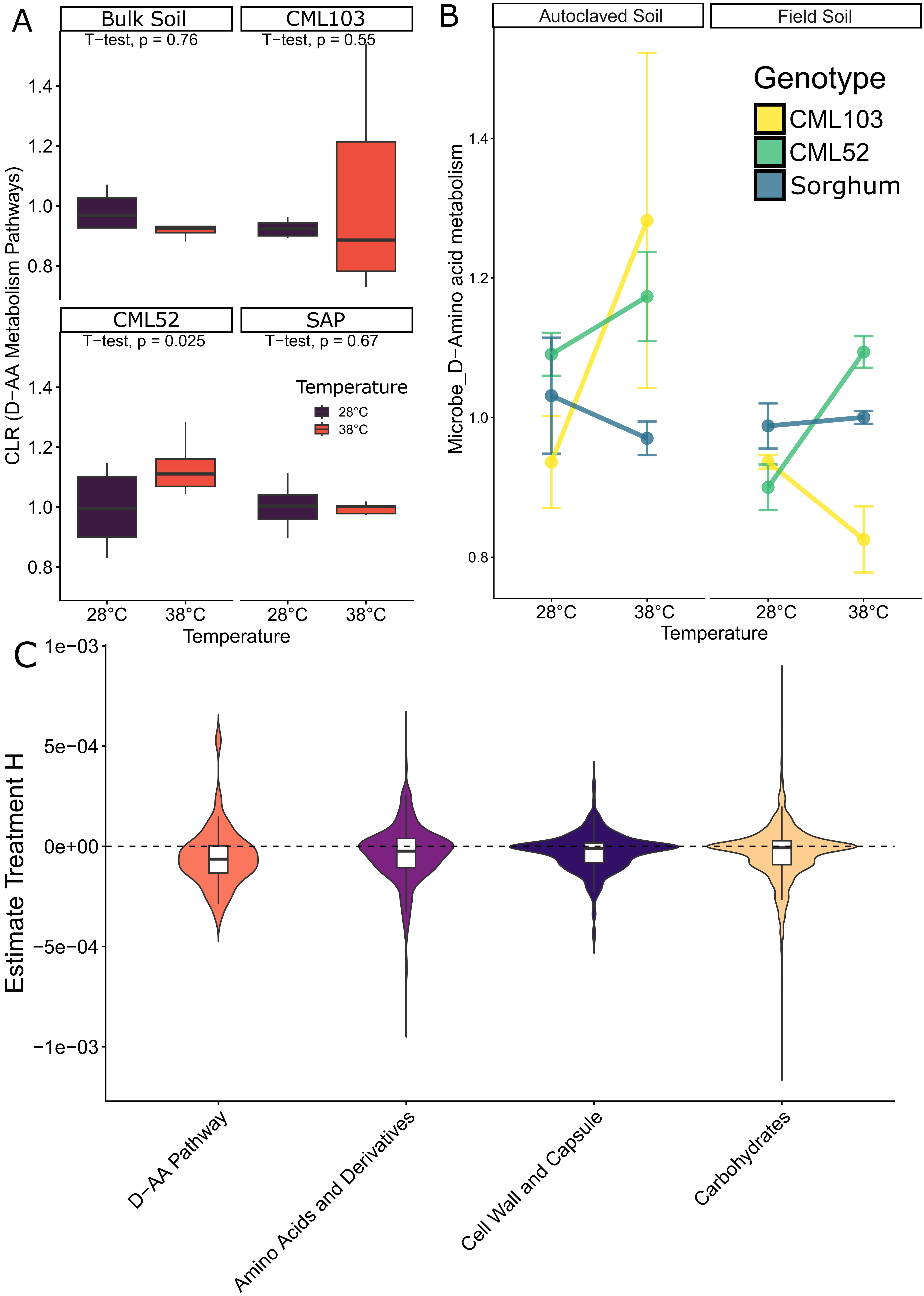
