## Supplementary material for "Investigating GERMs: How Genotype, Environment, and Rhizosphere Microbiome interactions underlie heat response in maize and sorghum": Descriptions of Supplementals

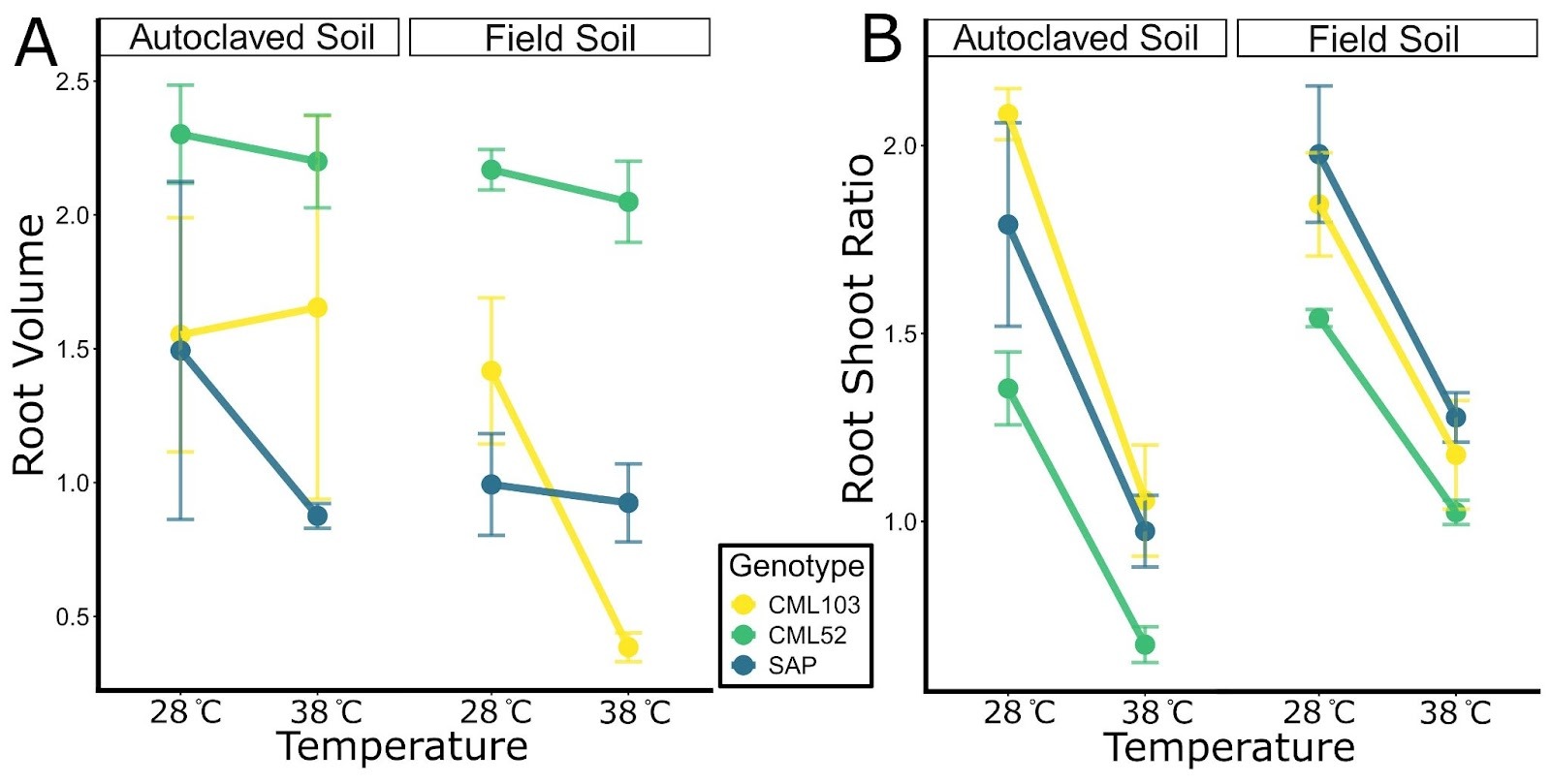

**Supplemental Figure 1. Additional plant phenotypes.** *Root volume was relatively stable in heat-tolerant plants and decreased rapidly in response to heat stress (A). Temperature had a consistent effect on the root-to-shoot ratio across soil treatments (B).*

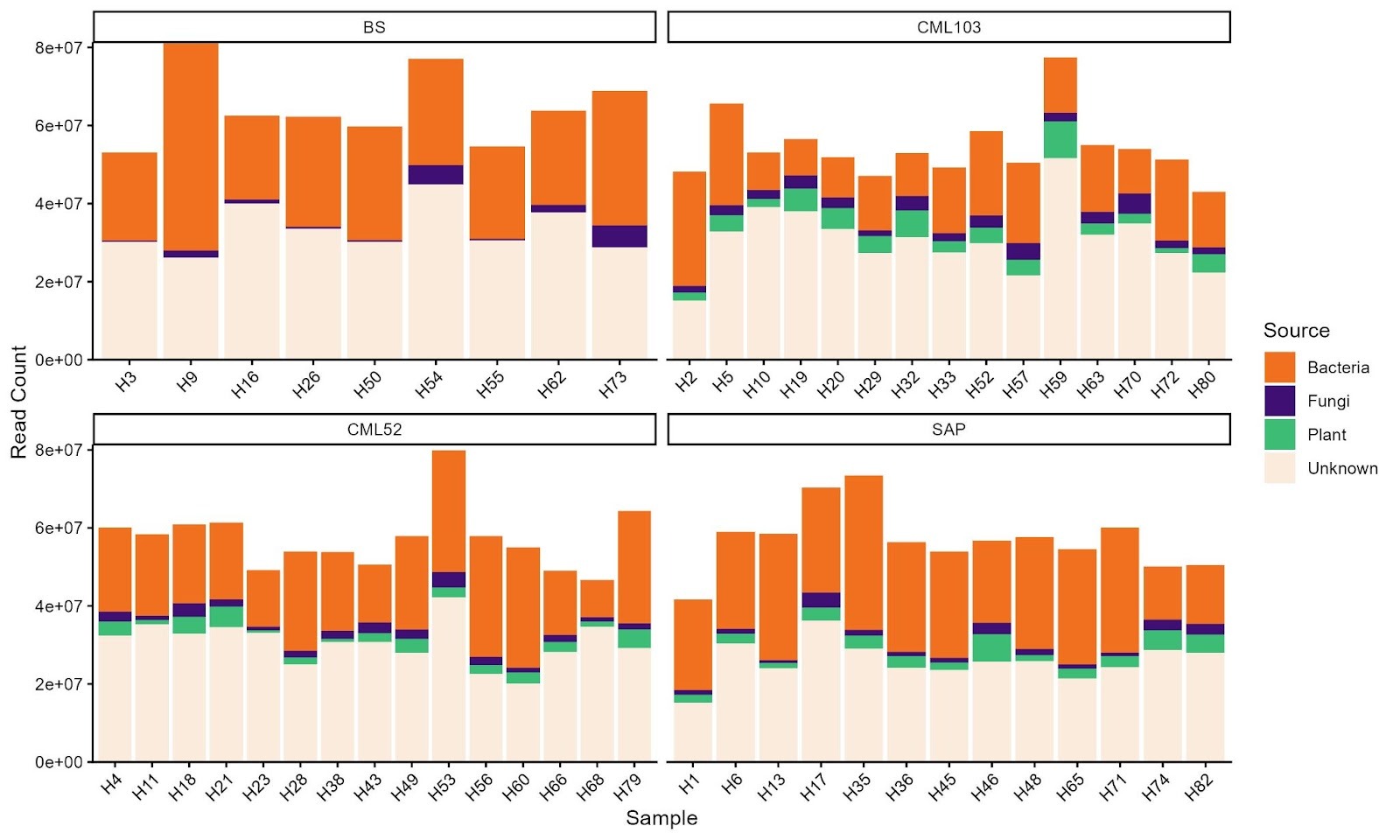

**Supplemental Figure 2. Proportion of reads assigned to plants and microbes.** *Read counts of all samples faceted by genotype and colored by assignment to plant, bacteria, or fungi.*

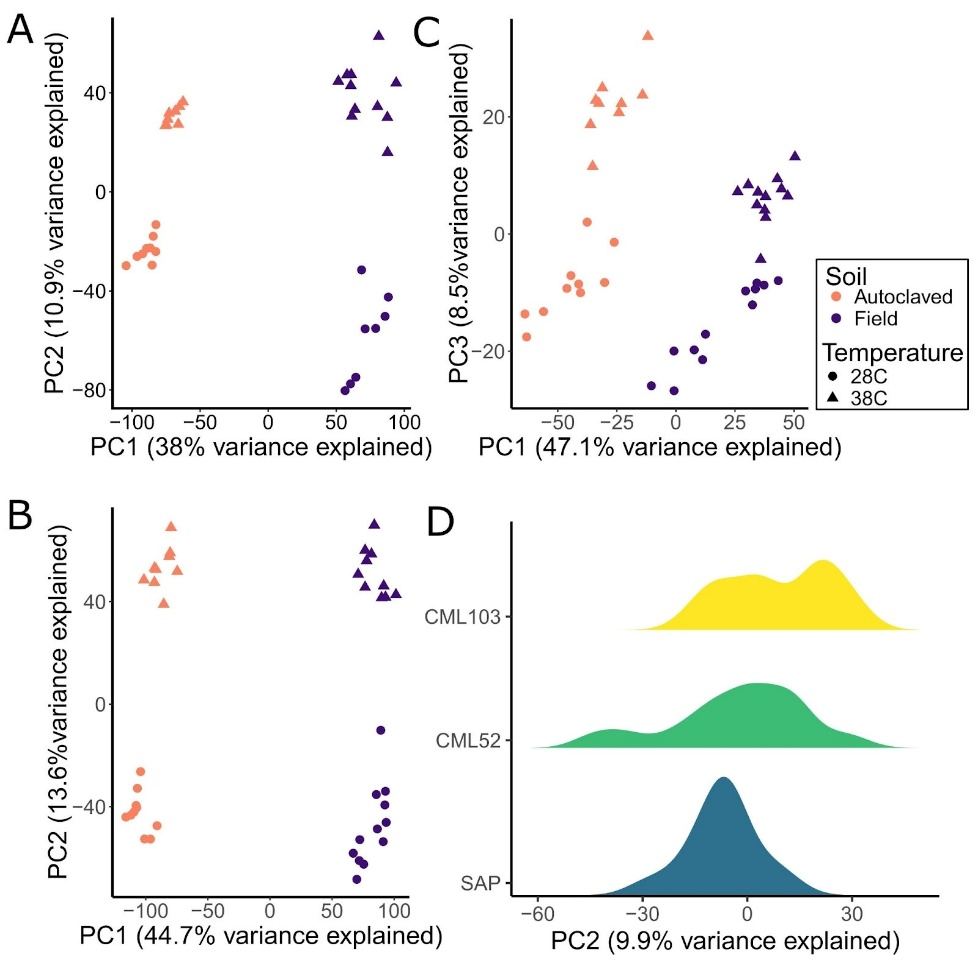

**Supplemental Figure 3. Beta-diversity analysis of community profiles from different sequencing types.** *Beta-diversity (Aitchison’s PCoA) of data from active 16S (A), all 16S (B), and transcriptomics (C), colored by soil type. PC3 is shown for transcriptomics, as PC2 is partially explained by genotype (D).*

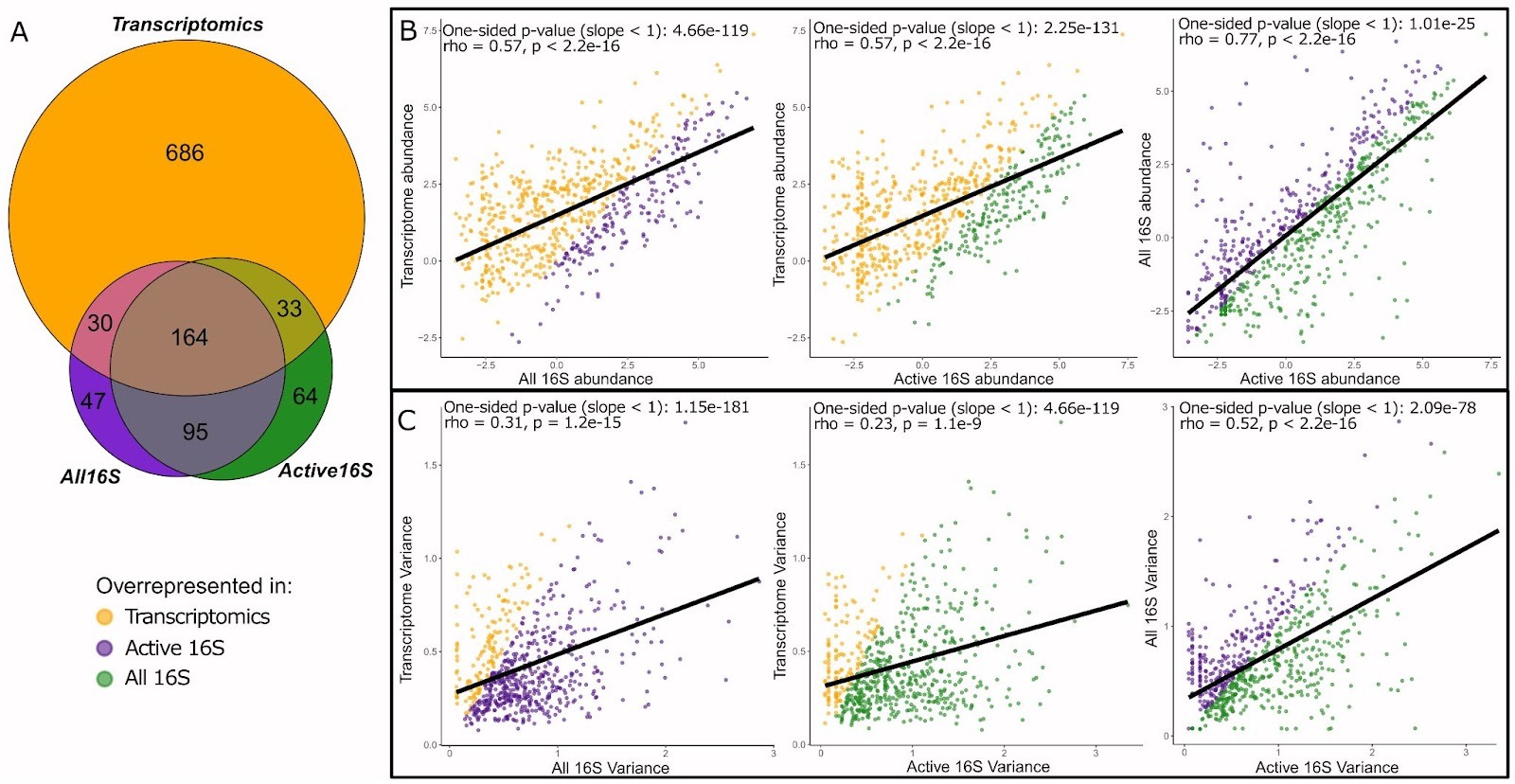

**Supplemental Figure 4. Comparison of sequencing methods.** *Venn diagram showing shared bacterial genera among sequencing types. Transcriptomics (orange) captured more unique genera than either all 16S (green) or active 16S (purple) (A). Correlation of abundance between sequencing methods, showing enhanced detection of low‐abundance taxa in transcriptomics (B). Variation between replicates, showing greater reproducibility in transcriptomic data (C). In all dot plots, the Spearman correlation trend lines are shown in black, and points are colored based on deviation from the expected 1:1 slope.*

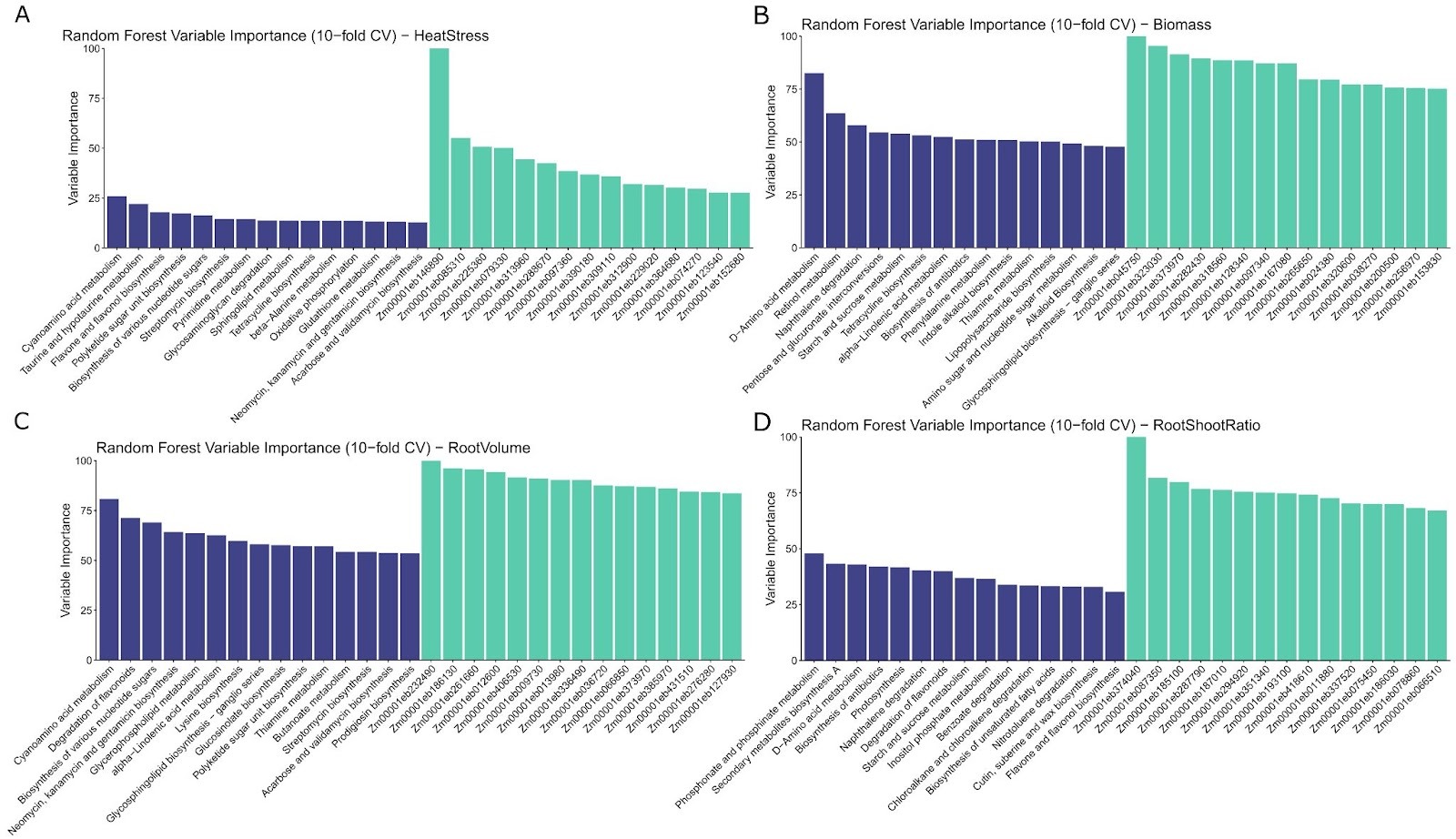
**Supplemental Figure 5. Random forest variable importance.** *The top 15 microbial pathways and maize genes identified by a random forest model for predicting biomass using 2,000 trees and 10-fold cross-validation for heat stress (A), biomass (B), root volume (C), and root-to-shoot ratio.*

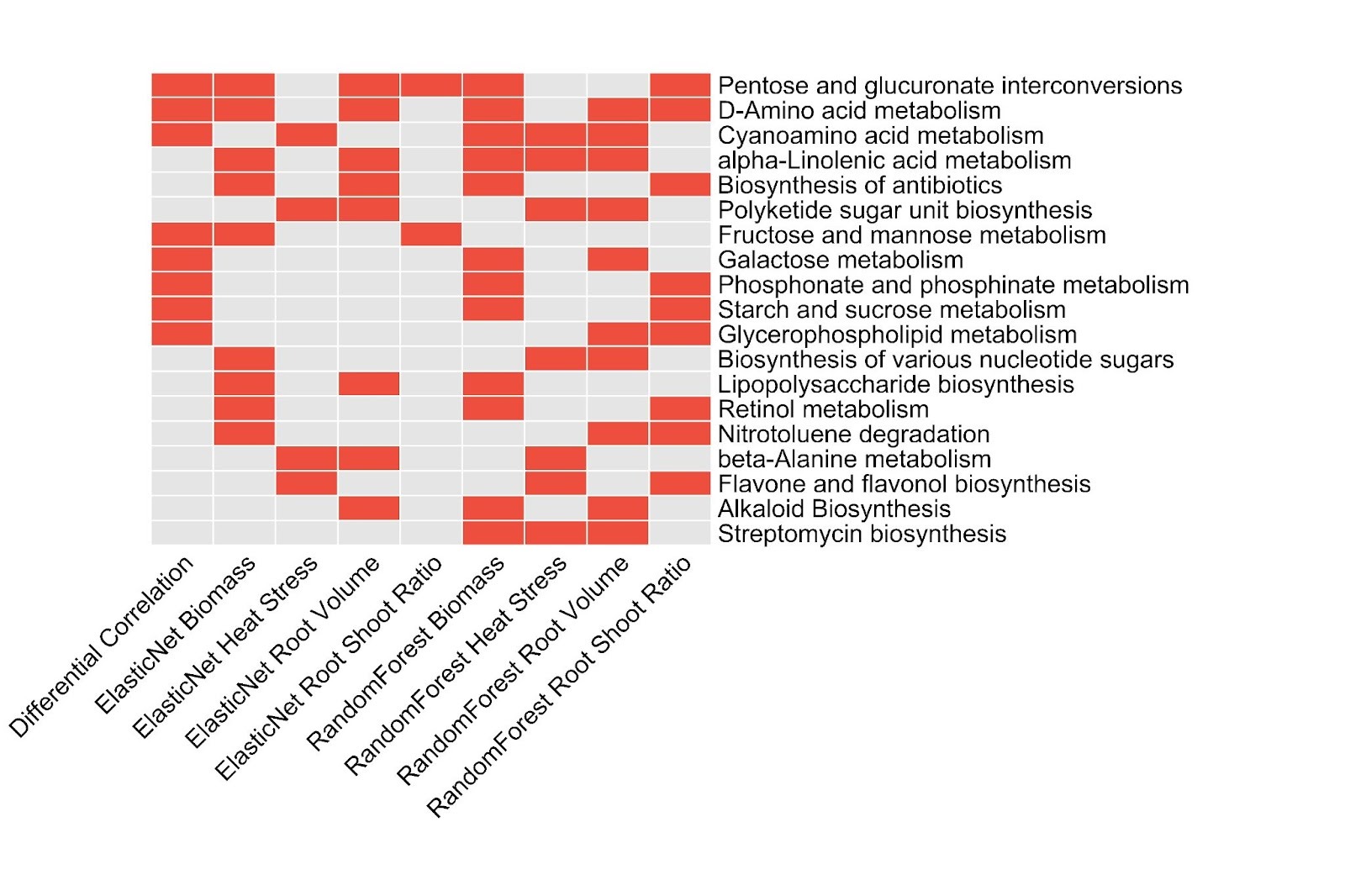

**Supplemental Figure 6. Candidate microbial mechanisms.** *Microbial Pathways present in at least three analyses of the differential bipartite correlation, Elastic Net, or the Random Forest, using biomass, heat stress, or root volume as the response variable.*

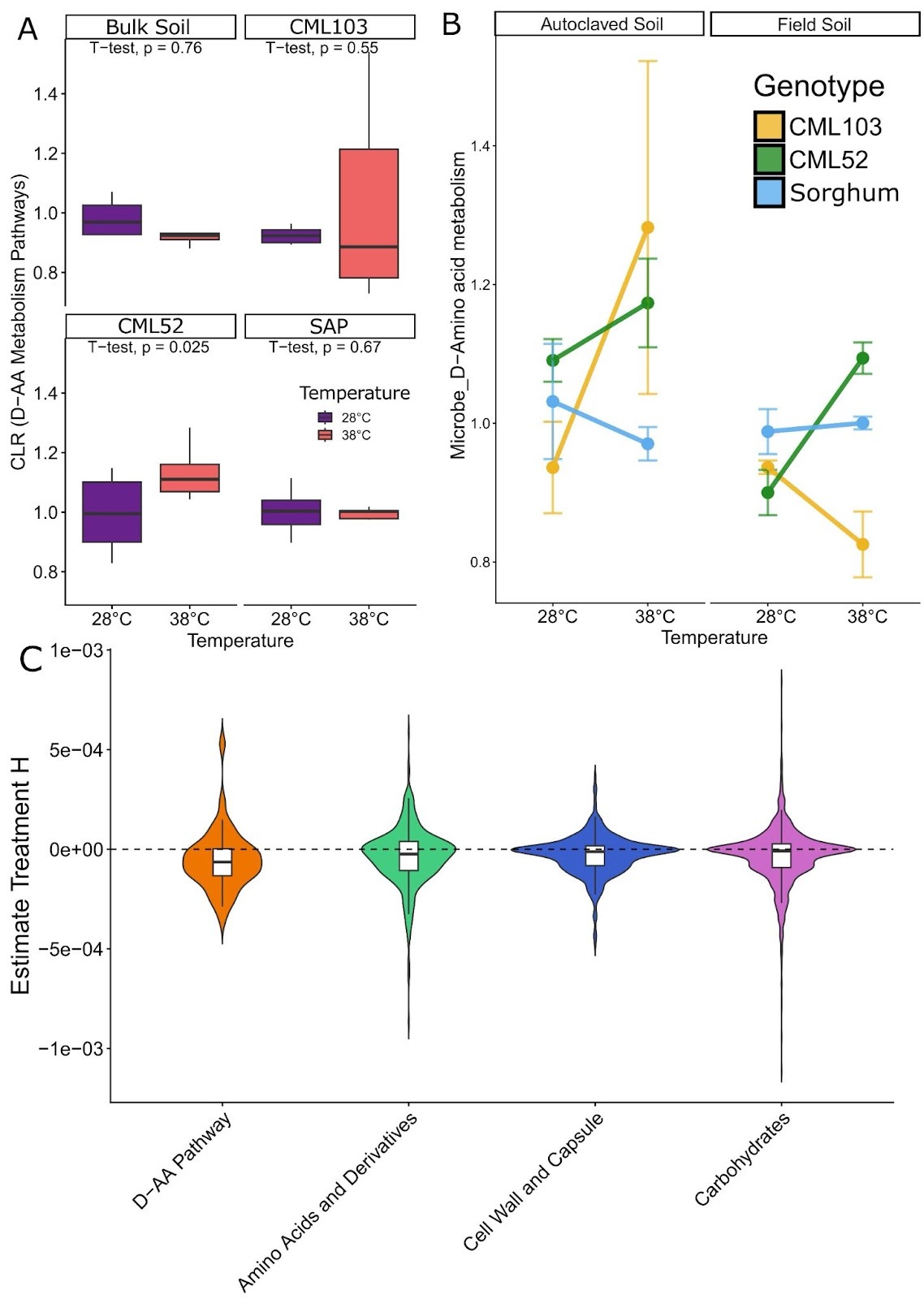

**Supplemental Figure 7. D-Amino acid metabolism across soils and genotypes.** *DAA metabolism was unchanged in bulk soil exposed to high temperatures and globally only increased in heat-tolerant CML52 (A). Soil type influenced DAA metabolism in a genotype-dependent manner (B). From publicly available data, heat stress on soil microbes in the absence of a plant results in decreased DAA metabolism (C).*

DataSets

SData1_DEmaize_sorghumgenes.xlsx

Differentially expressed genes from maize and sorghum across heat and control conditions, including log-fold changes, significance values, and orthology relationships used to compare conserved heat-responsive pathways across species. Also includes GO terms for genes identified in intersecting tests.

SData2_HeatDETranscriptomicsDatasets.xlsx

List of genes differentially expressed in high temperatures in maize and sorghum from this study and 5 additional publications, and the conversion of all geneID’s to maize, B3 v5.

Tables

ST1_SoilChemistry.xlsx

Soil chemistry profiles for field soil.

ST2_ExperimentalDesignMetaData.xlsx

Full metadata for all samples, including genotype, soil treatment, temperature condition, experimental block, and phenotypic measurements used in statistical and multivariate analyses.

ST3_ReadCounts.xlsx

Raw sequencing read counts for plant and microbial transcriptomes.

ST4_SorghumMaizeOrthogroups.xlsx

Orthogroup assignments linking maize and sorghum genes, enabling cross-species comparison of conserved genes.

ST5_CandidatePathways.xlsx

List of microbial pathways and plant genes identified as candidates in the integrated analyses—including correlations, elastic-net coefficients, and functional annotations—highlighting features most strongly associated with plant heat tolerance.
